# The NAT10 acetyltransferase modulates DNA damage-related factors and global 3D-genome architecture

**DOI:** 10.1101/2025.03.05.641614

**Authors:** Eva Bártová, Magdalena Skalníková, Lenka Stixová, Vlastimil Tichý, Filip Opálený, Jan Byška, Tomáš Brom, Soňa Legartová

## Abstract

We explored the role of NAT10 acetyltransferase in the DNA damage response, focusing on its impact on 3D-genome architecture and DNA repair proteins. Compared to NAT10 wild-type (wt), NAT10 deficiency reduced XPC, DDB2, and p53 protein levels. In TP53 double-null (dn) cells, the NAT10 protein was undetectable, and DDB2 was significantly down-regulated. Although NAT10 depletion caused DDB2 down-regulation, it did not affect the DNA repair functions of the DDB2 protein. To this fact, protein interaction analysis revealed that UVC exposure weakens the DDB2-p53 interaction while strengthening the bond between NAT10 and DDB2. Also, AlphaFold 3 prediction tools showed a more potent interaction between DDB2 and p53 than DDB2 and NAT10 proteins implying that NAT10 rather regulates the DDB2-p53 protein complex. These proteomic NAT10-dependent changes coincided with alterations in chromatin interactions, particularly in acrocentric chromosomes, studied by the Hi-C technique. However, 3D-genome rearrangement, caused by NAT10 deficiency and UVC irradiation, did not significantly impact post-translational histone modifications. Overall, NAT10 depletion alters the pool of key DNA repair proteins and induces substantial 3D-genome reorganization.

**Graphical abstract:** The effect of the NAT10 acetyltransferase depletion and UVC irradiation on 3D-genome nuclear architecture, histone signature, and the pool of selected DNA repair-related proteins. The figure was made using some icons adapted from BioRender software.

## Introduction

*NAT10* is a protein-coding gene that encodes the NAT10 protein belonging to the N-acetyltransferase superfamily that has acetyltransferase activity in the presence of acetyl coenzyme A (acetyl-CoA) and ATP (Ikeuchi, Kitahara et al., 2008). The function of the NAT10 protein is associated with various cellular processes, including RNA or protein acetylation and microtubule maintenance (Shen, Zheng et al., 2009). Acetylation is a co-transcriptional or post-translational modification where an acetyl group is added to distinct types of RNA or proteins influencing their function, stability, and localization within the cell nucleus and the cytoplasm (Lee, Hammaren et al., 2023). From this view, research on NAT10 revealed that this acetyltransferase is involved in cell cycle regulation, cellular proliferation, and response to DNA damage. For example, it is known that NAT10 is a specific RNA cytidine acetyltransferase catalyzing the formation of N(4)-acetylcytidine (ac4C) modification on mRNAs, 18S rRNA, and tRNAs (Arango, Sturgill et al., 2018, Ito, Horikawa et al., 2014, Sharma, Langhendries et al., 2015). Nuclear distribution studies showed that a high density of NAT10 is in the nucleolar compartment (Figure 1A), which documents its dominant role in the regulation of ribosomal RNA. Also, it is known that NAT10 is responsible for ac4C installation at position 1842 in 18S rRNA (Ito et al., 2014) and is essential for early cleavages of precursor rRNA at specific sites during 18S rRNA synthesis (Ito et al., 2014, Sharma et al., 2015) (Figure 1B). The structural analysis showed that NAT10 forms a symmetrical heart-shaped dimer (Figure 1C, Supplementary Figure 1A) (Zhou, Gamage et al., 2024). To this fact, it was revealed that ac4C modification in mRNAs strengthens mRNA stability and translation (Arango et al., 2018). Importantly, autoacetylation of NAT10 is required for its function which, among others, regulates transcription factors UBF1/2 that are responsible for rRNA synthesis (Cai, Liu et al., 2017). Although NAT10 occupies nucleoli and to a lesser extent the nucleoplasm, it was shown that the GFP-NAT10 deletion mutant (Δ989-1018) predominantly translocates into the cytoplasm with a weak signal in the nucleolus, while GFP-NAT10(Δ68-75) occupies both the nucleolus and the nucleoplasm (Tan, Zheng et al., 2018).

**Figure 1.**
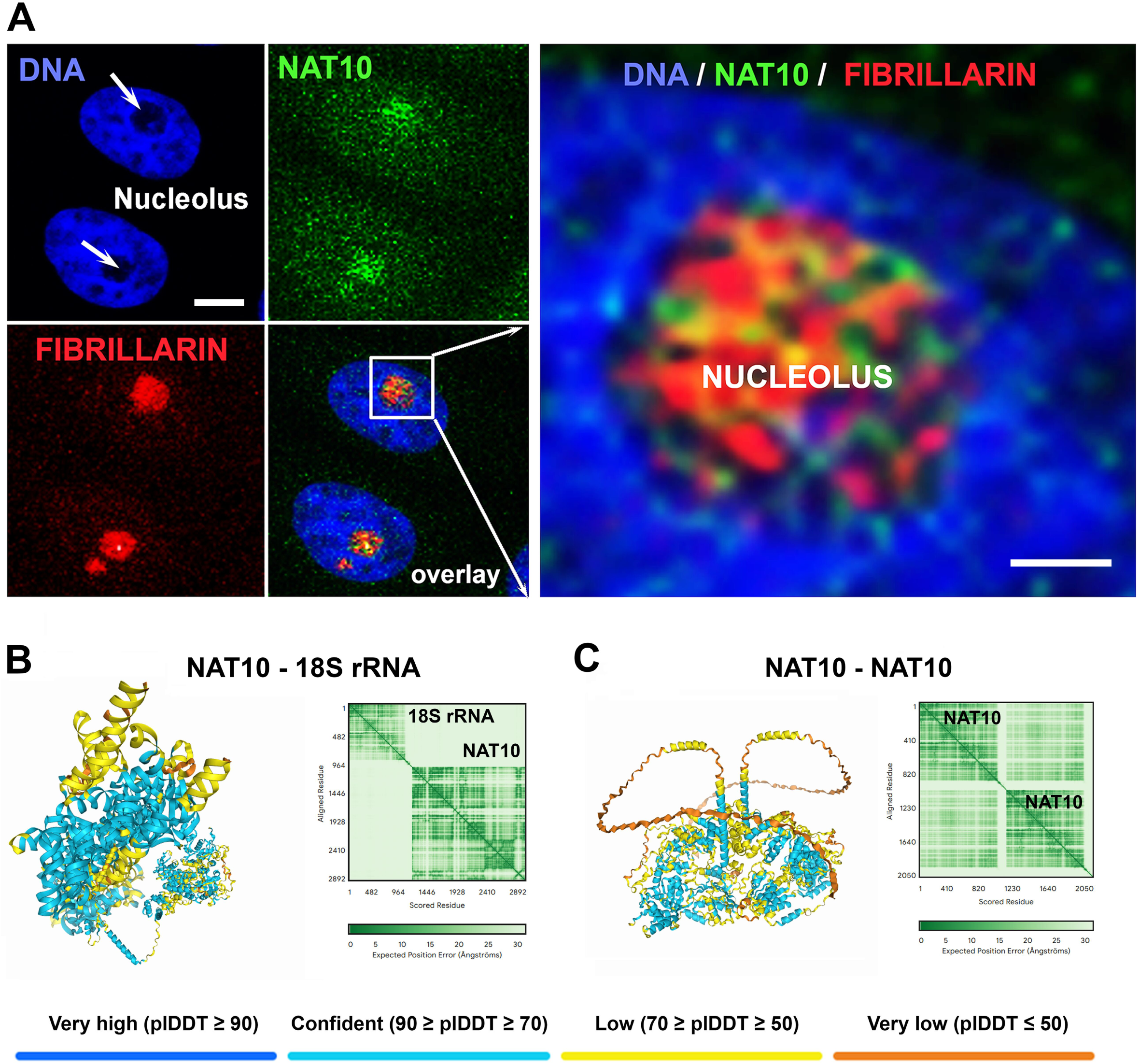
Nuclear localization of the NAT10 protein and the levels of selected DNA repair proteins in NAT10 wild-type and NAT10 double null cells. **(A)** The NAT10 acetyltransferase (green) occupies the nucleolus visualized by the antibody against fibrillarin (red). Nuclear DNA was stained by DAPI (blue). Bars represent 0.8 µm. **(B)** NAT10 interaction with 18S rRNA is documented with pTM = 0.51. **(C)** AlphaFold 3 model of NAT10 dimerization is shown (pTM = 0.54). Structures are colored using AlphaFold 3 pLDDT confidence score.

NAT10 can also mediate acetylation of lysine residues on histones, or p53 and MDM2 proteins, (Cai et al., 2017, Chi, Haller et al., 2007, Liu, Tan et al., 2016, Liu X 2018, Lv, Liu et al., 2003). It was shown that NAT10 stabilizes the p53 tumor suppressor by promoting MDM2 degradation under normal conditions and acetylates p53 to activate transcription of the p53 gene after DNA damage. It is well-known that p53 is a key player in the cellular response to DNA damage, and its acetylation status can influence its activity, which is important for coordinating the DNA damage response (Liu et al., 2016). These data document that NAT10 acetyltransferase may influence the acetylation status of specific proteins involved in DNA damage repair and cell cycle regulation. A very recent paper published by (Yang Z., 2023), shows that NAT10 plays a critical role in the repair of UVB-induced DNA damage lesions by regulating the expression of the key genes, including *DDB2*, which protein product is involved in the Nucleotide Excision Repair pathway (NER). These authors observed that the knockdown of *NAT10* enhanced the repair of UVB-induced DNA damage lesions via the stabilization of DDB2 mRNA. Also, these authors revealed that long-term UVB irradiation increases the NAT10 protein levels in mouse skin. On the other hand, we observed that the NAT10 function is not essential for the recruitment of ac4C RNA to UVA-microirradiated chromatin; thus, we suggested that another acetyltransferase could be responsible for the installation of ac4C in RNA appearing at UVA-damaged chromatin (Svobodova Kovarikova, Stixova, et al., 2023).

In this study, we introduce a new role of the NAT10 acetyltransferase in DNA damage response and the 3D-genome architecture studied in the HeLa cervix adenocarcinoma cell line. From this view, we analyzed the status of DNA damage repair-related proteins in non-irradiated and UVC-irradiated NAT10 wild-type (wt) and NAT10 double null (dn) cells. Also, we studied how p53 depletion affects the pool of the NAT10 protein and DNA repair proteins, including XPA, XPC, and DDB2 playing a role in the Nucleotide Excision Repair (NER) mechanism. Performing Hi-C analysis, providing data for chromatin interaction characteristics, especially Topologically Associated Domains (TADs) we concluded that the NAT10 depletion, and to a lesser extent irradiation by UVC light, causes significant rearrangement of the whole 3D-genome. TADs represent a genomic region where DNA sequences interact more frequently with each other than with sequences outside the domain. We especially showed that acrocentric chromosomes, forming the NAT10-abundant nucleoli, were rearranged. Also, we studied histone signature and observed that the level of specifically modified histones H3 and H4 remained relatively stable except for H3K9me3 reduced in NAT10 dn cells, and H3K79me1/me3 mostly diminished by UVC irradiation of this cell line. Together, despite the NAT10 protein being localized to and likely regulating specific compartments of the nucleoli and nucleolar processes, its depletion significantly alters the levels of non-nucleolar DNA repair-related factors and the global 3D-genome architecture.

## Results

### NAT10 deficiency reduced the level of the DDB2 and p53 proteins playing a role in the Nucleotide Excision Repair pathway

Here, we study the effect of NAT10 deficiency on DNA damage response. Using immunofluorescence combined with confocal microscopy we verified localization of the NAT10 acetyltransferase mainly in nucleoli (Figure 1A). Using western blots, in NAT10 wt and NAT10 dn cells, we analyzed levels of selected DNA repair proteins: p53, XPA, XPC, and DDB2 (Figures 2A-F). It is known that the DDB2 protein forms a heterodimeric complex with DDB1 which is essential in the NER process recognizing UV-damaged chromatin (Stoyanova, Roy et al., 2009, Tang & Chu, 2002, Wittschieben & Wood, 2003). Thus, we also studied potential multimeric interactions using the AlphaFold 3 prediction tool for the following proteins: NAT10, p53, DDB1, and DDB2 (Figure 2G-I and Supplementary Figure 1B-D, Supplementary Figure 2A and B).

**Figure 2.**
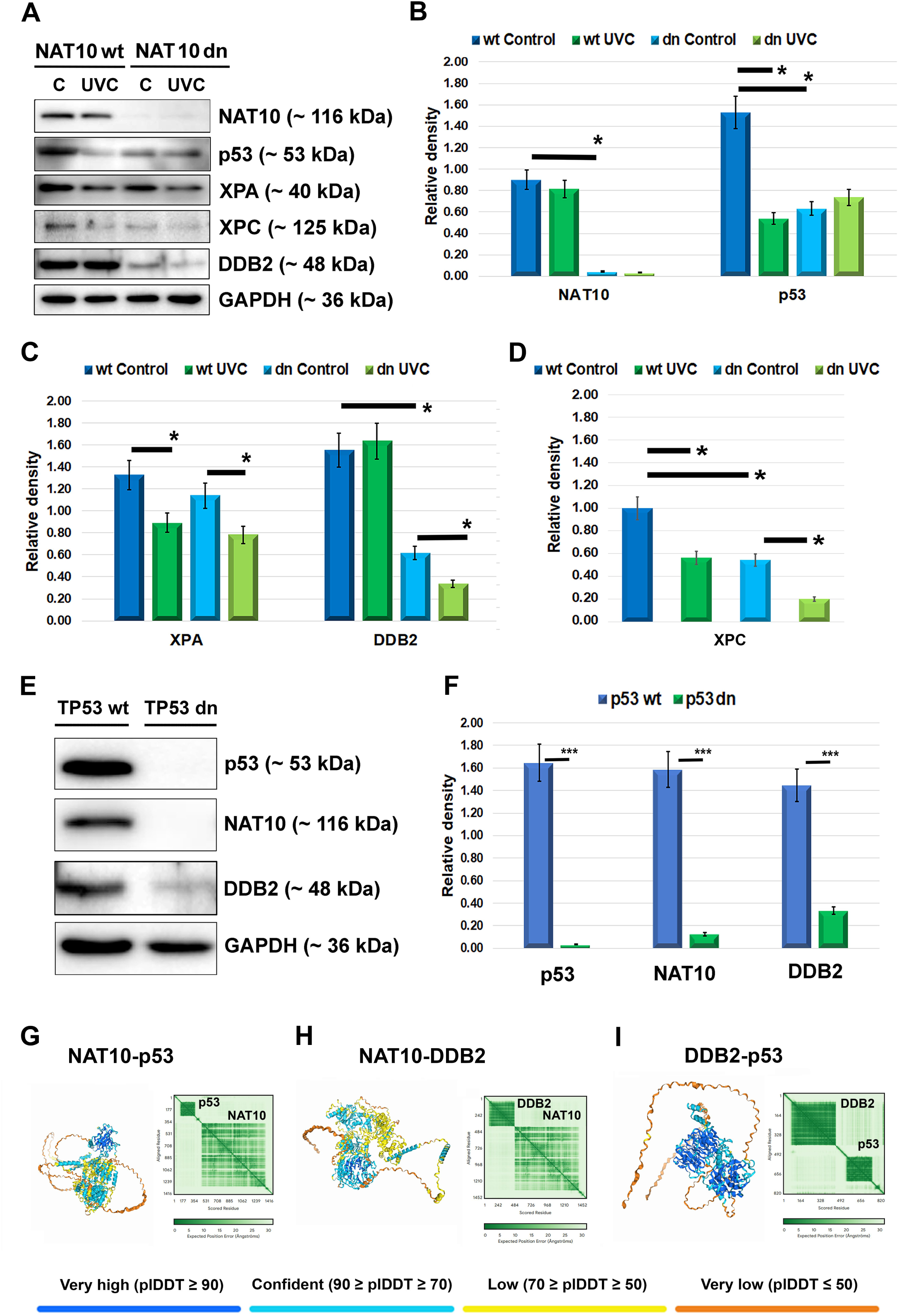
The levels of selected DNA repair proteins in NAT10 or TP53 wild-type and NAT10 or TP53 double null cells. **(A)** The following proteins were studied in non-irradiated and UVC-irradiated NAT10 wt and dn cells: NAT10, p53, XPA, XPC, and DDB2. In detail, panel (A) shows western blot analysis of proteins NAT10, p53, XPA, XPC, and DDB2 in non-irradiated and UVC-irradiated cells without and with a depletion of the NAT10 acetyltransferase. Data revealed lower p53, XPA, XPC, and DDB2 protein levels in NAT10 dn cells, with further reductions of XPA, XPC, and DDB2 protein levels upon UVC exposure. UVC irradiation also decreased the pool of p53, XPA, and XPC in NAT10 wt cells. Panels **(B-D)** show the quantification of western blot data from panel (A) using ImageJ software (NIH freeware, USA). Non-irradiated cells and cells irradiated by UVC light were analyzed. **(E)** TP53 deficiency led to a decrease in the pool of NAT10 and DDB2 proteins. Panel **(F)** shows the quantification of western blot data from panel (E) using ImageJ software (NIH freeware, USA). The protein levels were normalized to GAPDH (reference and loading control). The asterisks in panels B-D and F represent statistically significant differences with either a p-value ≤ 0.05 (*) or a p-value ≤ 0.001 (***). AlphaFold 3 models of (**G)** NAT10 interaction with p53 (pTM = 0.53), (**H**) NAT10 and DDB2 (pTM = 0.5), and (**I**) DDB2 and p53 protein (pTM = 0.47). Structures are colored with AlphaFold 3 pLDDT confidence score.

In these specifically designed experiments, we found using western blots a significant decrease in the pool of p53, XPA, XPC, and DDB2 proteins in non-irradiated NAT10 dn cells when compared with non-irradiated NAT10 wt counterparts (Figure 2A-D). Furthermore, a reduction in p53, XPA, and XPC proteins after UVC irradiation was observed in the NAT10 wt cell populations. Exposure to UVC light also diminished the XPA, XPC, and DDB2 protein levels in NAT10 dn cells, as shown in Figure 2A, C, and D.

From the view of the potential prediction of protein-protein interactions, AlphaFold 3 analysis suggests that p53 interacts with DDB2 and there is a weak interaction between NAT10-p53 and NAT10-DDB2 (Figure 2G-I). In comparison, NAT10-DDB2 interaction is potentiated in the presence of the p53 protein (Supplementary Figure 2A). Also, AIphaFold 3 confirmed a very strong interaction between DDB1 and DDB2 which influenced the complexation properties of NAT10 and DDB2 and p53 proteins (Supplementary Figure 2B).

### Interplay between p53, DDB2, and NAT10 in DNA damage response and repair

The p53 protein is also known to play a crucial role in DNA damage response (Williams & Schumacher, 2016, Zhang, Liu et al., 2011). This protein can regulate the expression of various genes involved in DNA repair mechanisms, including DDB2, summarized in Capuozzo, Santorsola et al. (Capuozzo, Santorsola et al., 2022). The p53 protein can induce the expression of the *DDB2* gene by binding to its promoter, which in turn influences the activity of the DDB1-DDB2 complex which is crucial for the NER mechanism (Scrima, Konickova et al., 2008). As mentioned above, this complex specifically recognizes UV-induced DNA damage, which is then repaired to maintain genomic integrity (Stoyanova et al., 2009). In this study, we described, not only by AlphaFold 3, a mutual relationship between p53, DDB2, and NAT10 proteins (Figure 2G-I). Also, in TP53 wild-type cells (p53 wt) we observed high levels of DDB2 and NAT10 proteins, while in TP53 double null cells (p53 dn) we found a barely detectable pool of these proteins studied (Figure 2E and F). Especially, the level of the NAT10 acetyltransferase was affected by the TP53 deficiency (Figure 2E and F).

### Depletion of NAT10 abrogates an interaction between p53 and DDB2 and UVC irradiation changes interactions between NAT10 and DDB2

Using the Proximity Ligation Assay (Figure 3A), we showed that in NAT10 dn cells the strength of interaction between p53 and DDB2 was reduced in comparison to NAT10 wt counterpart, while UVC irradiation did not change interaction properties between p53 and DDB2 proteins in both NAT10 wt and NAT10 dn cells (Figure 3B-E). Also, in NAT10 wt cells, we observed that UVC radiation significantly strengthened interaction properties between NAT10 and DDB2 proteins (Figure 3F). As mentioned above, AlphaFold 3 analysis showed that p53, in a complex with NAT10 and DDB2, might strengthen an interaction between NAT10 and DDB2 (compare Figure 2H with Supplementary Figure 2A).

**Figure 3.**
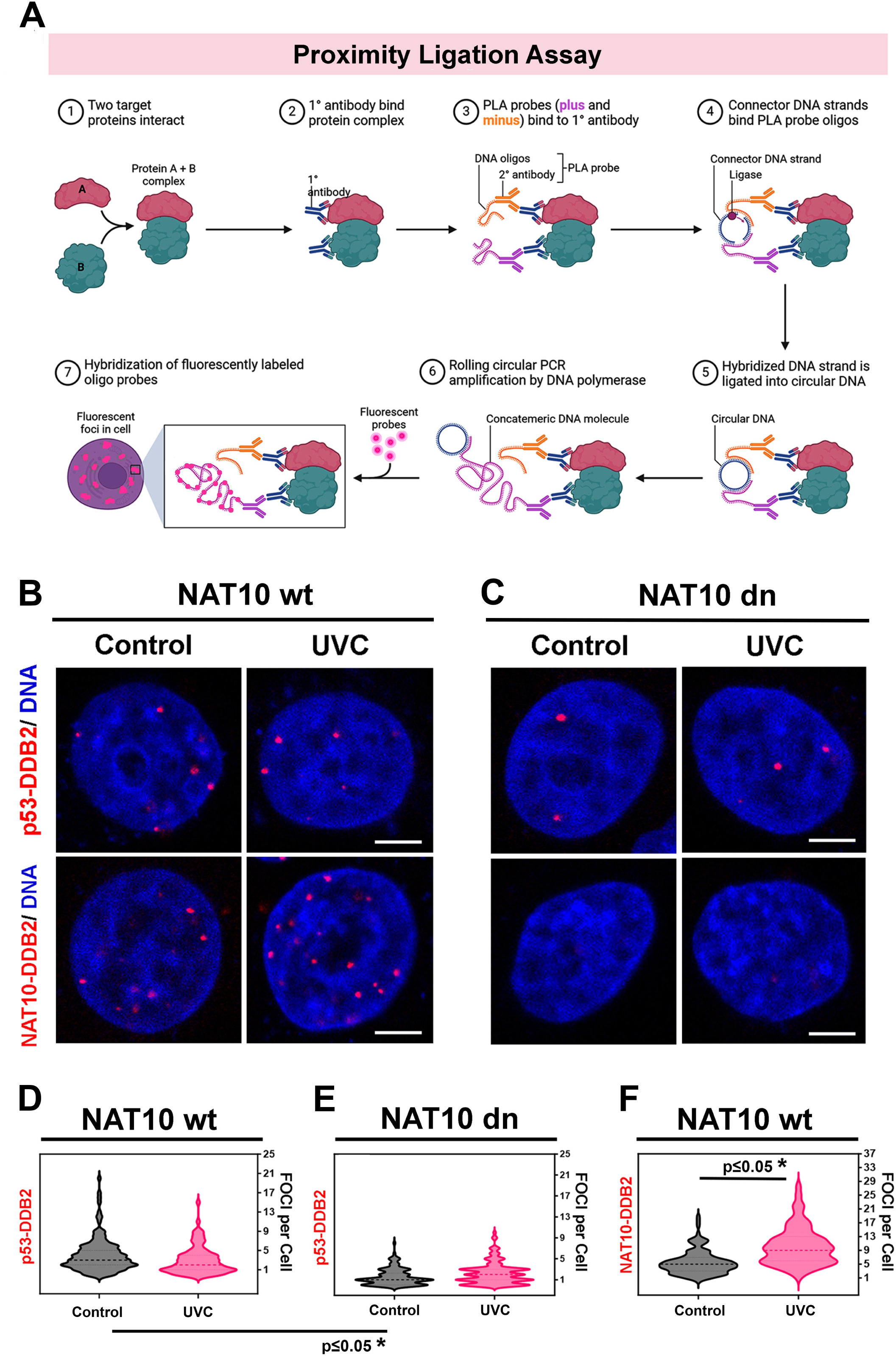
A Proximity Ligation Assay (PLA) showing p53-DDB2 and NAT10-DDB2 interaction properties. Interaction properties between p53-DDB2 and NAT10-DDB2 in human NAT10 wt and NAT10 dn cells were studied using PLA. **(A)** The panel shows an example of PLA methodology, adapted from BioRender software. In **(B)** NAT10 wt and **(C)** NAT10 dn cells, after fixation, the PLA was performed using antibodies to detect the p53-DDB2 or NAT10-DDB2 complexes in both non-irradiated and UVC-irradiated NAT10 wt and NAT10 dn cells. Red dots indicate amplified interaction signals, with cell nuclei counterstained with DAPI, used for visualization of DNA content. NAT10-DDB2 in NAT10 dn acted as a control, showing no external impact on PLA outcomes. This section quantifies PLA signals (dots per nucleus) for **(D)** the p53-DDB2 complex in non-irradiated and UVC-irradiated NAT10 wt cells, **(E)** p53-DDB2 in non-irradiated and UVC-irradiated NAT10 dn cells. **(F)** The UVC irradiation significantly increased NAT10-DDB2 PLA signals in NAT10 wt cells, suggesting enhanced interaction. Non-irradiated cells and cells irradiated by UVC light were analyzed. We used ImageJ software (NIH freeware, USA) to analyze the number of PLA signals. The statistical analysis was performed using GraphPad Prism 9 software (USA) and the nonparametric Mann-Whitney *U*-test. The asterisk in panel F represents statistically significant differences with a p-value ≤ 0.05 (*).

Based on this result we continued to study the nuclear distribution of the DDB2 protein in NAT10 wt and NAT10 dn cells, and we have observed tiny DDB2-positive foci that appear in DAPI-dense nuclear region (heterochromatin), DAPI-poor region of the genome (euchromatin), and inside the compartment of nucleoli (Figure 4A-D). In the following experiments, we tested the hypothesis of whether NAT10 depletion influences the recruitment of the DDB2 protein to microirradiated chromatin. We observed identical recruitment of the DDB2 protein to UVA-microirradiated genomic regions in both NAT10 wt and NAT10 dn cells (Figure 4E). It means that even though the fact that NAT10 depletion caused DDB2 down-regulation (Figure 2A and C), this change did not affect the functional properties of the DDB2 protein in DNA damage repair (Figure 4E).

**Figure 4.**
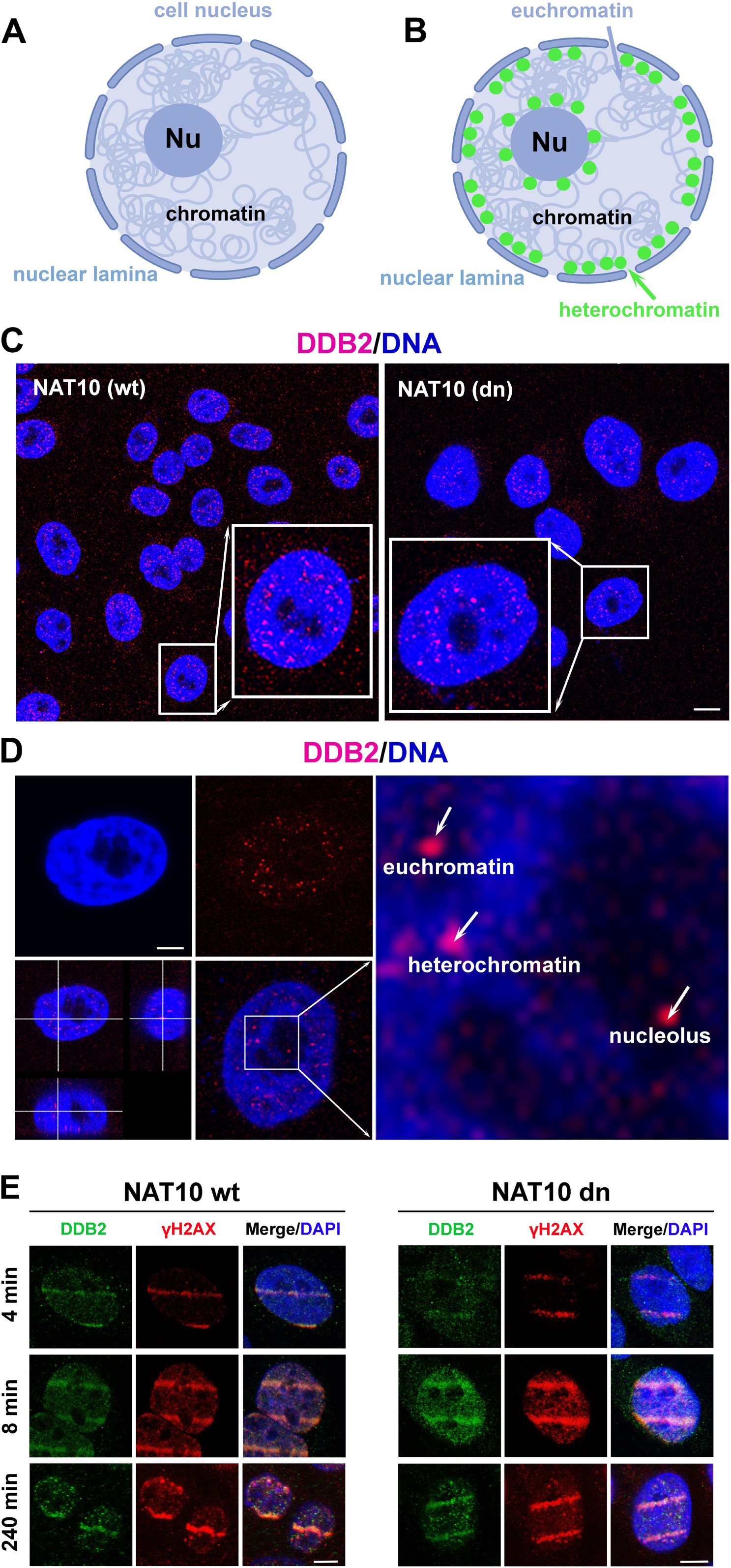
Nuclear distribution profile of the DDB2 protein in NAT10 wt and NAT10 dn cells. **(A, B)** Animation documents chromatin density inside the cell nucleus (see graphical illustration created by BioRender software). Panel **(A)** shows cell nucleolus (Nu) and nuclear lamina in dark blue, and panel **(B)** shows localization of heterochromatin (green) decorating the periphery of cell nucleus and nucleolus, while euchromatin (pale blue) is shown as de-condensed treads inside the cell nucleus. An analysis of the DDB2 nuclear distribution pattern was done using immunohistochemistry in **(C)** NAT10 wt and NAT10 dn cells. **(D)** The NAT10 protein (red) was accumulated into tiny, well-visible foci in DAPI-dense (considered as heterochromatin), DAPI-poor (euchromatin, more decondensed) genomic region as well as inside compartment of nucleoli. Panel **(E)** shows representative images of the DDB2 protein recruitment to UVA-microirradiated chromatin in NAT10 wt and NAT10 dn cells. Microirradiation using a 405 nm laser line was performed, followed by immunofluorescent staining for DDB2 and DNA damage marker γH2A.X. DAPI was used for nuclear counterstaining in blue fluorescence. The scale bar represents 7.5 μm.

### NAT10 depletion and UVC irradiation affect the 3D-genome organization

We used the Hi-C method to identify the physical interactions between different regions of the genome, providing insights into how the genome is organized within the nucleus and how this organization influences gene expression and cellular function. We inspected all human autosomes and chromosome X in human cervical adenocarcinoma HeLa cells, and we noticed that the 3D-genome nuclear organization is changed in both NAT10-depleted and UVC-irradiated cells. Focusing on acrocentric chromosomes carrying the nucleolus organizing regions (NOR), we also observed interaction changes when we compared the chromatin of NAT10 wt and NAT10 dn cells or these cells exposed to UVC light (Figures 5-7 and Supplementary Figures 3-5). First, we present differential loop analysis showing distinctions in the unspecific loops and conditional specific loops of all human chromosomes. For instance, differential loop analysis shows distinctions between the aggregated features of common (unspecific loops) and unique chromatin loops (conditional specific loops) in different samples. In this case, we compared two groups in the following experimental setups: (A) chromatin of NAT10 wt non-irradiated cells with those in NAT10 dn non-irradiated cells (Figure 5A); (B) chromatin of NAT10 wt non-irradiated cells was compared with NAT10 wt UVC-irradiated cells (Figure 5B); or we studied 3D-genome changes in (C) NAT10 dn non-irradiated cells when compared with NAT10 dn UVC-irradiated cells (Figure 5C). The heatmap uses colors to indicate the interaction frequency at chromatin loop anchors. The red region shows interactions with highly enriched properties, while the pale blue regions show a low degree of chromatin loop interactions. The bottom-left red region in the heatmaps suggests an asymmetrical interaction pattern, where contacts might extend from the loop anchor to adjacent genomic areas. This could be indicative of an extended enhancer region interacting with multiple target genes or one-sided interactions that do not fully bring both loop anchors together. The central red region in the heatmaps indicates strong looping interactions directly between the loop anchor regions, which are symmetrically enriched. This is typical for loops that are stable and form a closed chromatin conformation, such as those that bring together specific enhancers and promoters to activate gene transcription. In detail, the low differences in enrichment scores (ES) between the two experimental conditions show the non-significant changes in chromatin structure (compare NAT10 wt / NAT10 dn or NAT10 wt / NAT10 wt UVC or NAT10 dn / NAT10 dn UVC in left panels of Figure 5A-C). Conversely, the highest differences in ES between the two experimental conditions mean a new formation of unique chromatin loops (compare NAT10 wt / NAT10 dn or NAT10 wt / NAT10 wt UVC or NAT10 dn / NAT10 dn UVC in middle and right panels of Figure 5A-C).

**Figure 5.**
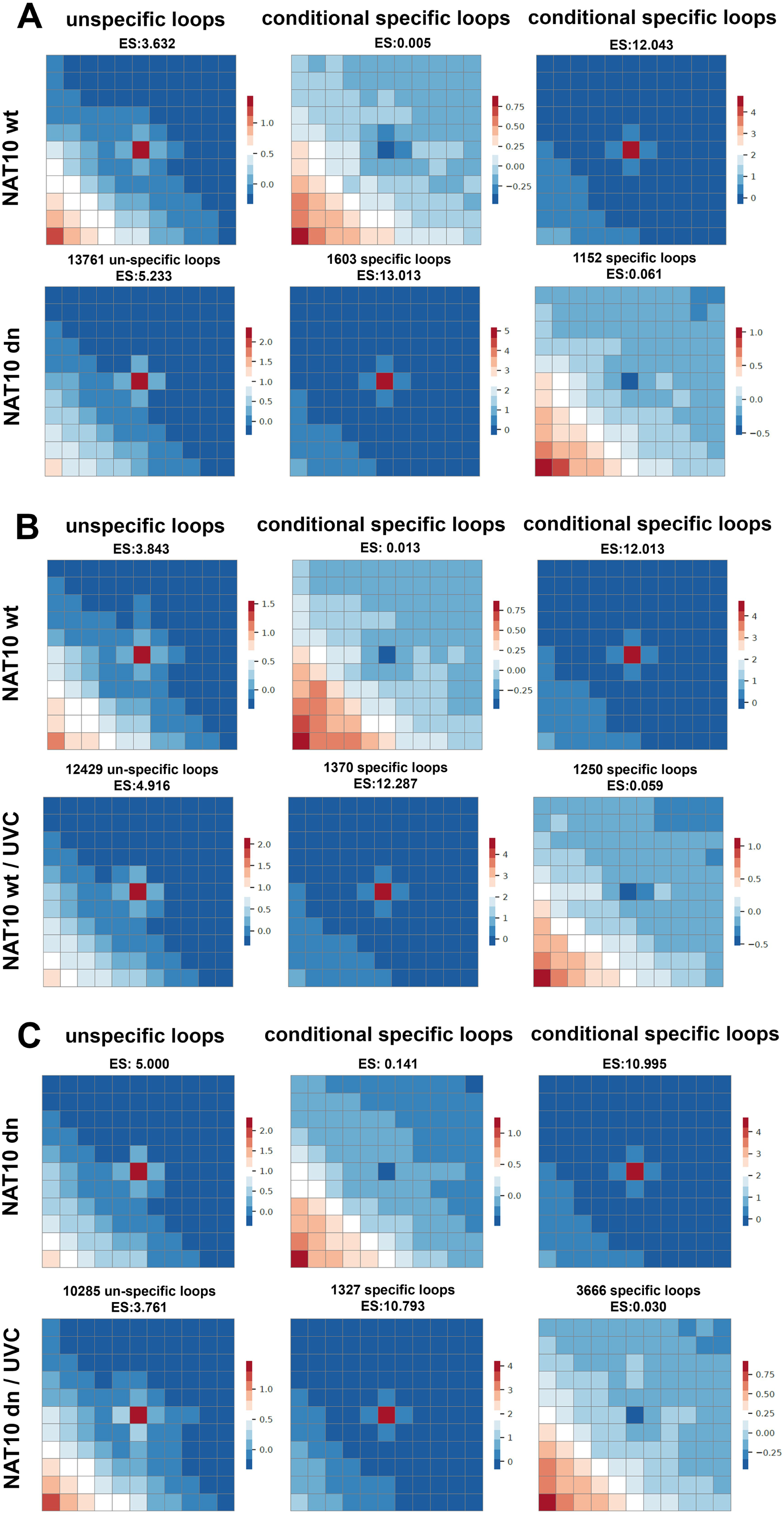
Changes in 3D-genome nuclear architecture of human chromosomes analyzed in non-irradiated and UVC-irradiated NAT10 wt and NAT10 dn cells. The results of the differential loop analysis are shown for the following comparisons: **(A)** NAT10 wild-type non-irradiated cells versus NAT10 double null non-irradiated cells; **(B)** NAT10 wild-type non-irradiated cells versus NAT10 wt UVC-irradiated cells; **(C)** NAT10 double null non-irradiated cells versus NAT10 double null UVC-irradiated cells. The contact maps show the differential loop analysis for common (unspecific loops) (left), and unique (conditional specific) loops (the middle showing an increased number of loops and the right panels showing a reduced number of loops) in the comparison between two experimental conditions as shown in **(A)**, **(B)**, and **(C)**. Labeling ES means an enrichment score. The color scale indicates ES, with red regions representing higher loop enrichment and blue regions representing loops with reduced interaction properties.

Also, the number of reduced and highly enriched chromatin loops are displayed in volcano plots documented in Figure 6A-C showing 1603 highly enriched loops and 1152 loops with reduced interaction properties in NAT10 dn non-irradiated cells (Figure 6A). Similarly, 1370 highly enriched loops and 1250 loops with reduced interactions in NAT10 wt UVC-irradiated cells (Figure 6B). And finally, 1327 highly enriched loops and 3666 loops with reduced interaction properties in NAT10 dn UVC-irradiated cells (Figure 6C). In this study, we also show contact maps of the 3D-organization of human acrocentric chromosomes (Supplementary Figures 3-5). An example of pronounced differences is documented in Figures 7A and B showing contact maps of human acrocentric chromosome 13. In this case, we observed the most pronounced differences in 3D-nuclear architecture, especially in the region around 50-70 Mbp of HSA13 when we compared Hi-C maps of (a) NAT10 wt / NAT10 wt UVC with (b) NAT10 wt / NAT10 dn cells (Figure 7A, B). Data suggests that UVC irradiation affects whole HSA13, while NAT10 deficiency selectively affects specific regions of HSA13. For an explanation, red-marked regions in contact maps (on the right side of the panel) indicate a higher number of interactions compared to the control, while blue-marked ones represent a lower number of interactions. Results of our analysis indicate that both NAT10 depletion and UVC irradiation significantly changed the whole 3D-genome organization, which is also reflected in the 3D-rearrangement of all human acrocentric chromosomes (Figure 7C).

**Figure 6.**
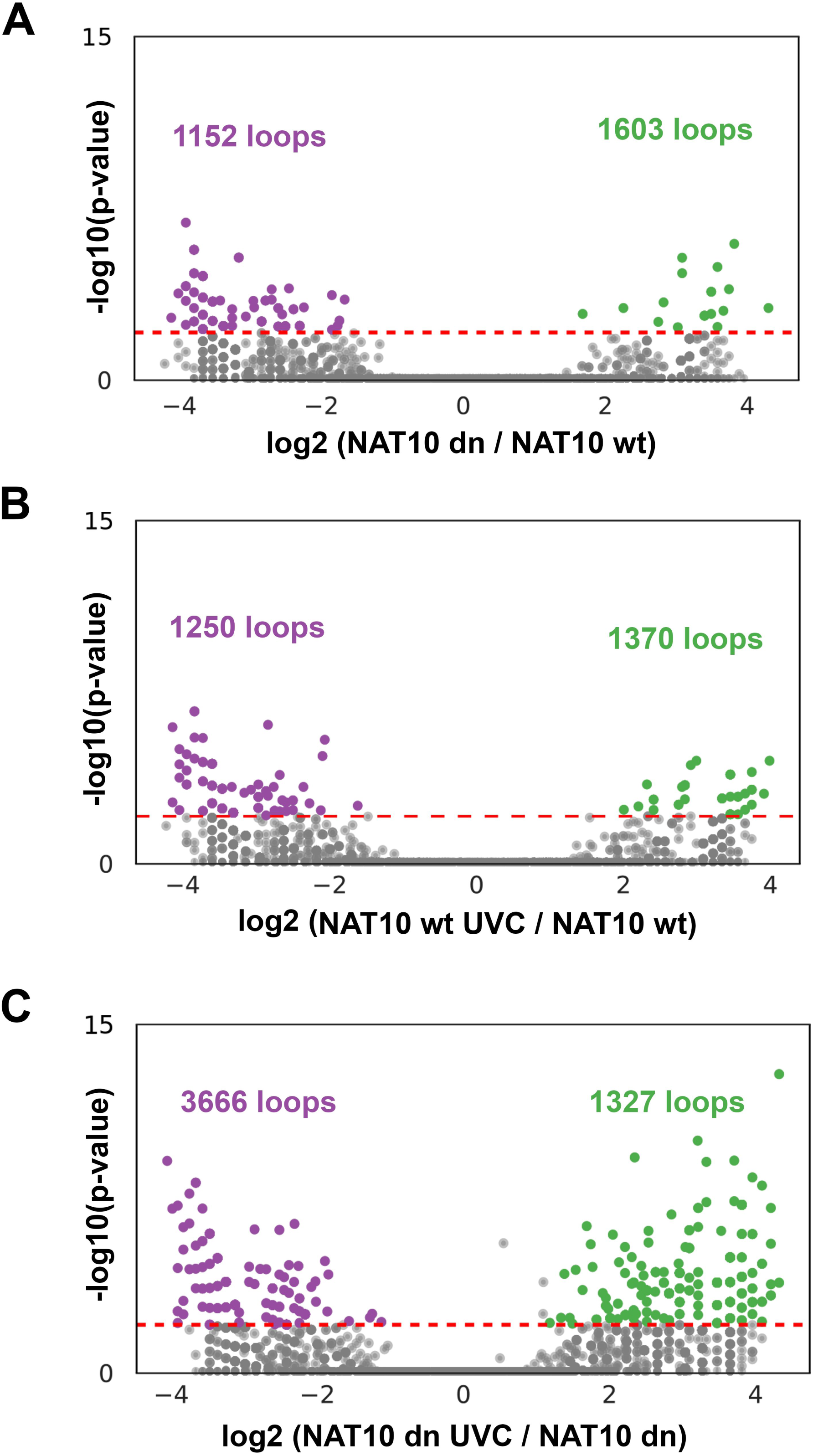
The number of reduced and highly enriched chromatin loops in NAT10 wt and NAT10 dn non-irradiated and UVC-irradiated cells. Volcano plots illustrate the results of the differential analysis of chromatin loops (cLoops). These plots depict the relationship between fold change and statistical significance (p-value = 0.01, red line) of individual loops under two experimental conditions. The x-axis represents the logarithmic values of the fold change, indicating the degree of change in interactions between the two conditions. The y-axis shows the negative logarithmic value of the p-value (-log10(p-value)), representing the measure of statistical significance. Differentially enriched chromatin loops are highlighted in green (highly enriched) and purple indicates a reduced number of loops. Insignificantly changed loops are shown in grey. The number of chromatin loop interactions was studied in **(A)** NAT10 dn non-irradiated cells when compared with NAT10 wt counterpart, **(B)** NAT10 wt UVC-irradiated cells compared with NAT10 wt non-irradiated cells, and **(C)** NAT10 dn UVC-irradiated cells compared with NAT10 dn non-irradiated cells.

**Figure 7.**
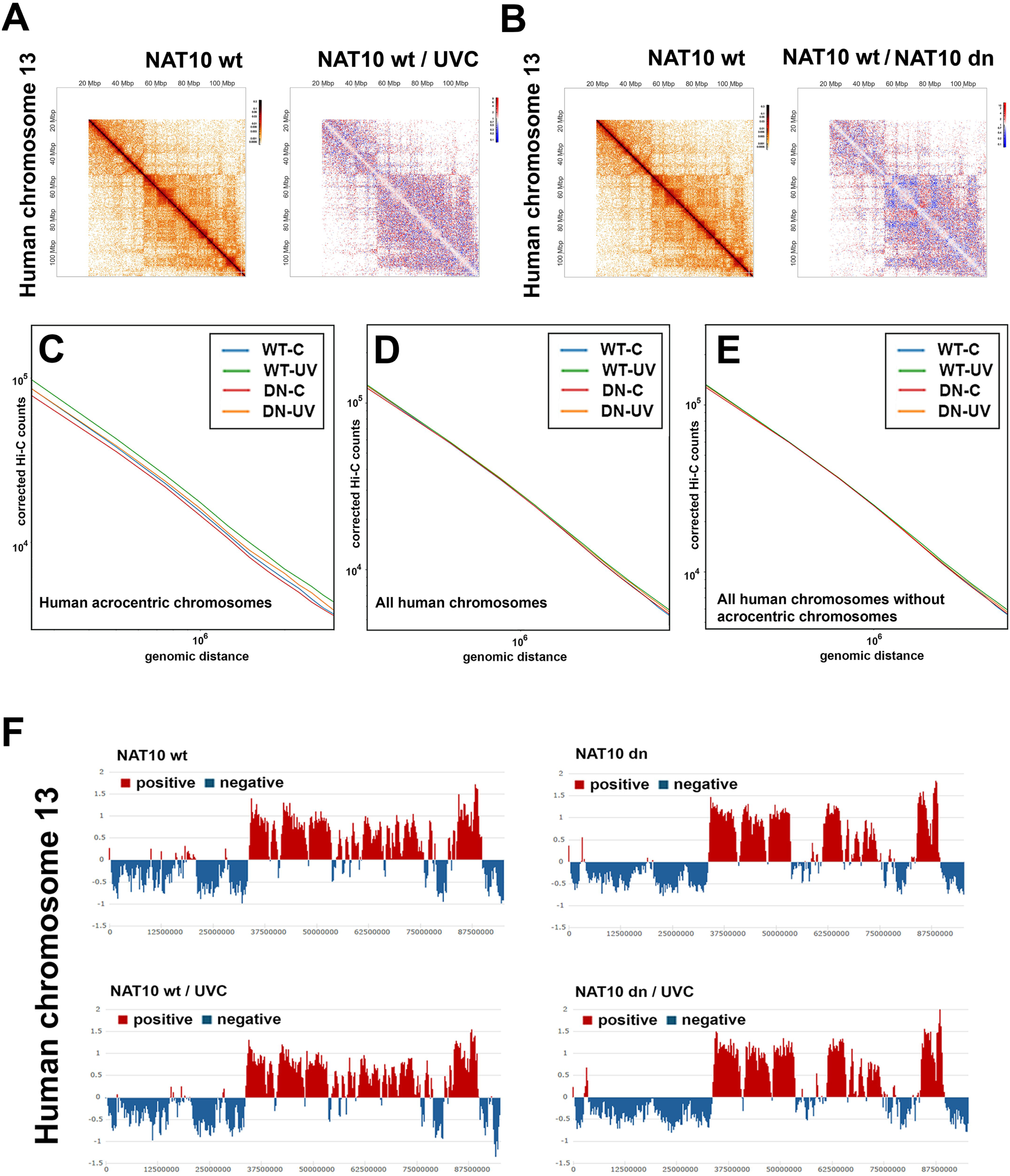
An example of changes in the 3D-genome nuclear architecture of human acrocentric chromosomes is caused by NAT10 deficiency and UVC irradiation. Changes in the 3D-organization (see contact maps and A/B compartments) of human acrocentric chromosome 13 (HSA13) were studied in non-irradiated and UVC-irradiated NAT10 wt and NAT10 dn HeLa cells. Genome-wide contact maps are shown in panels **(A, B)** for acrocentric chromosome HSA13 in 360 kb per bin. Red-blue panels show difference maps of acrocentric chromosome 13 visualized using the HiGlass software studied in non-irradiated NAT10 wt and NAT10 wt/UVC-irradiated cells or NAT10 wt and NAT10 dn cells. The missing data in panels A and B are shown in white color. Hi-C data were also analyzed as an average over **(C)** all acrocentric chromosomes and compared with the average of **(D)** all human chromosomes and **(E)** all chromosomes without acrocentric ones. Panel **(F)** shows compartment switching regions (B-to-A) (A in red shows active and open, positive, chromatin, and B in blue shows condensed and inactive, negative, chromatin) of human chromosome 13 analyzed in non-irradiated and UVC-irradiated NAT10 wt and NAT10 dn HeLa cells. The A/B compartments were identified according to eigenvector analysis, at a medium resolution of 250,000 base pairs. Decomposition from normalized correlation maps was applied.

In this case, the Hi-C data examining the number of interactions related to genomic distances were averaged over all acrocentric chromosomes (Figure 7C) and compared with the averages of all human chromosomes (Figure 7D) and all human chromosomes excluding the acrocentric ones (Figure 7E). This analysis showed the most pronounced NAT10 and UVC radiation-dependent chromatin changes in acrocentric chromosomes (Figure 7C). UVC irradiation increases the number of chromatin loop interactions, while NAT10 depletion decreases the number of interactions suggesting an opposite effect on chromatin structure in the case of acrocentric chromosomes. On the other hand, chromatin loop interactions were affected to a lesser extent by experimental setups in all human chromosomes (Figure 7D and E). For explanation, there were more significant differences among experimental setups when we studied acrocentric chromosomes and compared them with all human chromosomes and all human chromosomes without acrocentric ones (Figure 7C-E).

Also, we studied compartment-switching regions (B-to-A) for human chromosome 13 analyzed in non-irradiated and UVC-irradiated NAT10 wt and NAT10 dn HeLa cells. The analysis documented changes in A/B compartments, especially caused by NAT10 depletion. The most significant differences were found in the region 5×10^7^-7.5×10^7^ bp when we compared NAT10 wt and NAT10 dn cells (Figure 7F).

To specify the effect of NAT10 depletion and UVC irradiation on chromatin interaction properties, we show an example of TADs for human chromosome 13 (Figure 8A-F, Supplementary Figure 6, and Supplementary Table 1). This data shows significant differences in TADs when comparing NAT10 wt with NAT10 wt UVC-irradiated cells (Figure 8A and B), NAT10 wt with NAT10 dn cells (Figure 8C and D), and NAT10 dn with NAT10 dn UVC-irradiated cells (Figure 8E and F). The more pronounced changes we observed in NAT10 deficient cells when compared with their UVC-irradiated counterpart (see especially Supplementary Table 1).

**Figure 8.**
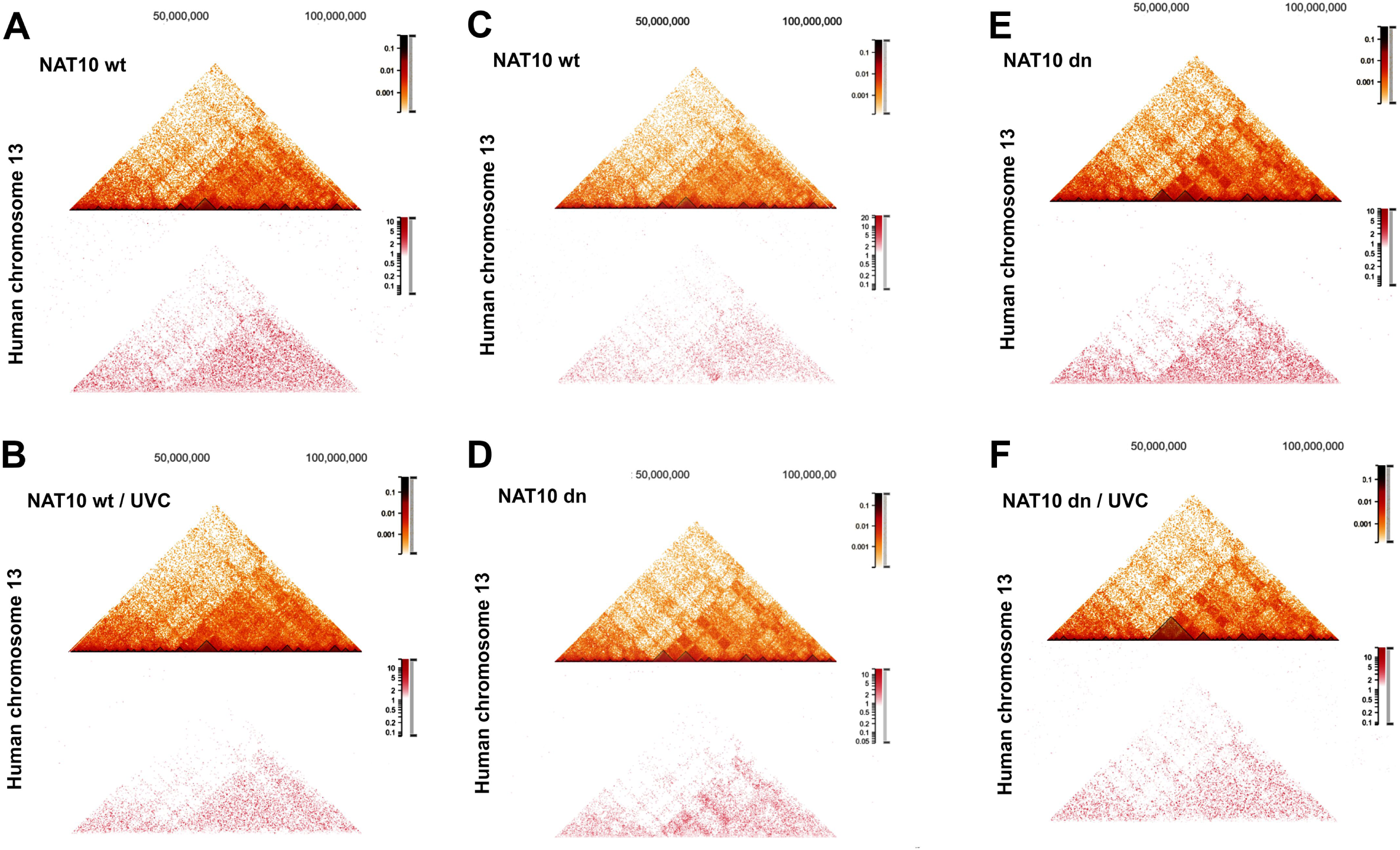
An example of changes in TADs of human acrocentric chromosome 13 is caused by NAT10 deficiency and UVC irradiation. Changes in TADs (shown in resolution 320 kB per bin) of HSA13 were studied in non-irradiated and UVC-irradiated NAT10 wt and NAT10 dn HeLa cells. Hi-C contact maps are shown with respective TAD annotations for all experimental events (upper orange panel in A-F). A difference map (bottom red panels) highlighting regions where NAT10 wt has more interaction than NAT10 wt/UVC **(A)**, where NAT10 wt/UVC has more interaction than NAT10 wt **(B)**, where NAT10 wt has more interaction than NAT10 dn **(C)**, where NAT10 dn has more interaction than NAT10 wt **(D)**, where NAT10 dn has more interaction than NAT10 dn/UVC **(E)**, and where NAT10 dn/UVC has more interaction than NAT10 dn **(F)**.

### Depletion of the NAT10 acetyltransferase and UVC-irradiation affect histone signature

Because chromatin is not only DNA but also histones, here, we continue to study nuclear architecture in a more complex way. In parallel with changes in chromatin interactions and TADs, caused by NAT10 depletion and UVC radiation, we analyzed changes in histone signature. Using western blots, we detected the level of the following histone markers, specific for both euchromatin and heterochromatin (Figure 9A and B): H3K9me1/me2/me3, H3K36me3, H3K79me1/me2/me3, H3K9ac, and H4ac. Analysis was performed in non-irradiated and UVC-irradiated NAT10 wt and NAT10 dn cells (Figure 9C). We observed that UVC light significantly reduced levels of H3K9me2 or H3K79me3 in UVC-irradiated NAT10 dn cells when compared with non-irradiated counterparts (Figure 9C, D, and E). In both NAT10 wt and NAT10 dn cells, UVC irradiation reduced H3K79me1 (Figure 9C and E). The level of H3K9me3 was decreased in both NAT10-depleted cells and NAT10 wt cells exposed to UVC irradiation (Figure 9C and D). Surprisingly, the global level of acetylation markers on histones H3 and H4 as well as H3K36me3 were stable in non-irradiated and UVC-irradiated NAT10 wt and NAT10 dn cells (Figure 9C-F).

**Figure 9.**
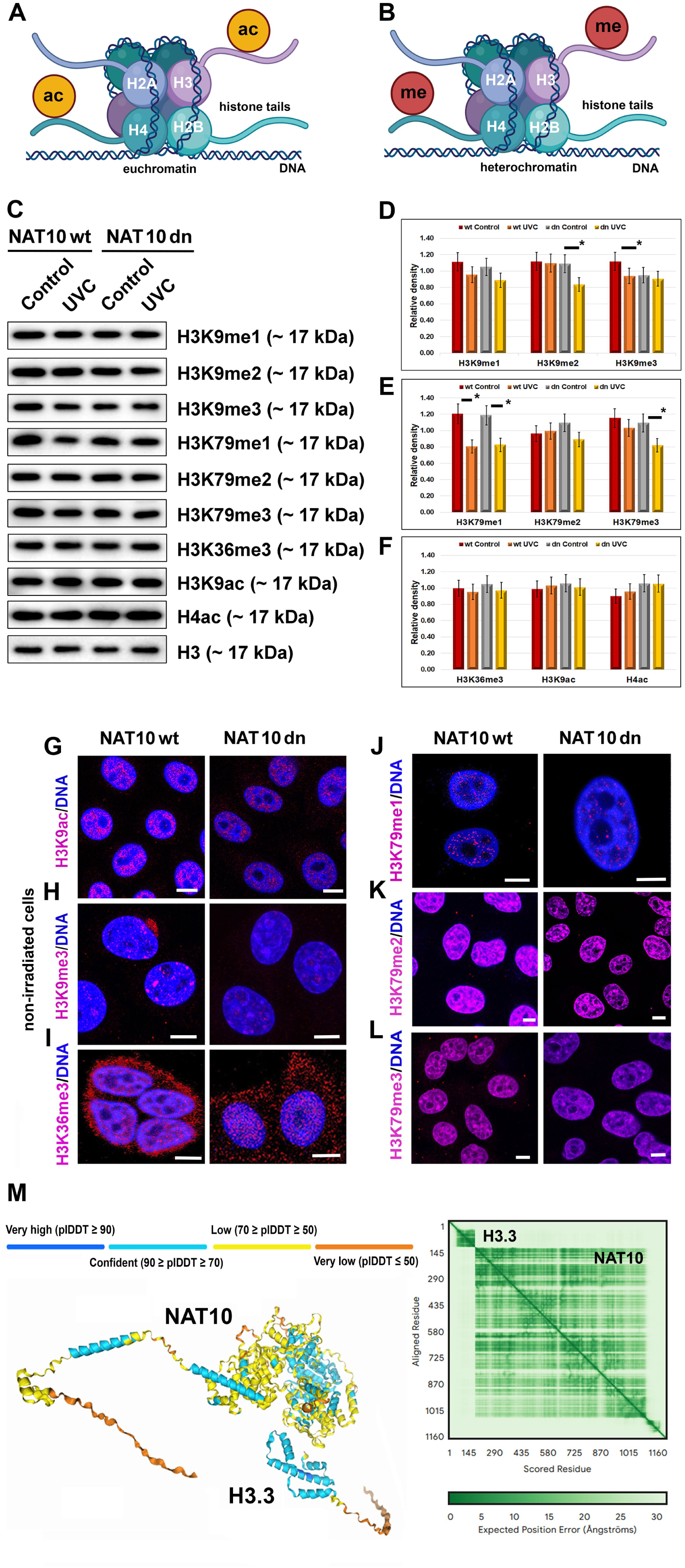
The level of post-translationally modified histones in non-irradiated and UVC-irradiated NAT10 wt and NAT10 dn cells. The levels of selected histone markers, characterized for **(A)** euchromatin or **(B)** heterochromatin, were studied in non-irradiated and UVC-irradiated NAT10 wt and NAT10 dn cells. Analyses were performed using western blots. **(C)** In non-irradiated and UVC-irradiated NAT10 wt and dn cells, the level of the following histone markers was analyzed: H4ac, H3K9ac, H3K9me1/me2/me3, H3K36me3, and H3K79me1/me2/me3. **(D-F)** Western blot data were quantified by the ImageJ software. Statistically significant differences at p ≤ 0.05 are shown by asterisk (*). Using immunofluorescence and confocal microscopy, the following histone markers were analyzed in NAT10 wt and NAT10 dn cells: **(G)** H3K9ac, **(H)** H3K9me3, **(I)** H3K36me3, **(J)** H3K79me3, **(K)** H3K79me2, and **(L)** H3K79me3. AlphaFold 3 predicted a low degree of interaction between the NAT10 protein and human core histone H3.3C (pTM = 0.57). Structures are colored with AlphaFold 3 pLDDT confidence score.

Also, we used quantitative immunofluorescence and confocal microscopy to study the nuclear distribution of H3K9ac, H3K9me3, H3K36me3, and H3K79me1/me2/me3 (Figure 9G-L). In NAT10 wt and NAT10 dn cells, we observed subtle changes in nuclear density and distribution of the above-mentioned histone marks. Both observations, by western blots and immunofluorescence, showed only subtle changes in histone signature caused by NAT10 depletion and UVC irradiation. This conclusion was also confirmed by the AlphaFold 3 prediction tool showing no interaction between NAT10 and the variant of histone H3 (H3.3C) (Figure 9M). Taken together, we show that despite the immense regulatory effect of the NAT10 depletion of 3D-genome architecture, the NAT10 acetyltransferase has no role in the regulation of histone signature.

## Discussion

Acetyltransferases play vital roles in the post-translational modification of proteins, both histone and non-histone, as well as RNAs, regulating diverse biological processes such as cell metabolism, gene expression, and cellular signaling. In this study, we identified a key role for the NAT10 acetyltransferase in maintaining the integrity of the DNA damage response (DDR). Specifically, we observed that the depletion of NAT10 leads to the downregulation of DNA damage-binding protein 2 (DDB2), as shown in Figure 2A and C. This relationship between NAT10 and DDB2 was previously reported (von Eyss, Maaskola et al., 2012, Yang Z., 2023) where NAT10 knockdown was found to enhance the repair of UVB-induced DNA lesions by stabilizing DDB2 mRNA. However, direct evidence of a functional interaction between NAT10 and DDB2 proteins during DNA damage repair has not been demonstrated until now. Here, we provide new insights, showing that NAT10 depletion directly impacts the DDB2 protein pool and that UVC irradiation strengthens the interaction between NAT10 and DDB2 (Figure 3F). To verify the mentioned observations, we ran AlphaFold 3 predictions and we confirmed the dimerization of NAT10, which seems to be important for the acetyltransferase activity of this protein, as also discussed by (Zhou et al., 2024). Also, AlphaFold 3 analysis showed that the p53 protein, in a complex with NAT10 and DDB2, potentiates an interaction between NAT10 and DDB2. Thus, we suggest that the NAT10-DDB2-p53 protein complex might be functionally more important in DNA damage response than heterodimer NAT10-DDB2 (compare Figure 2H with Supplementary Figure 2A; see left-bottom square in AlphaFold 3 heat maps). These results imply a novel interplay between NAT10 and the tumor suppressor protein p53. To support this claim, in NAT10-deficient cells, we observed reduced p53 levels, and in p53-depleted cells, NAT10 was under the detection level of western blots (Figure 2A and B). This interdependence was extended to DDB2, as p53 depletion also led to its downregulation (Figure 2E and F). Although p53 and NAT10 are known to function in distinct pathways (p53 primarily regulates cell cycle control and DNA repair, and NAT10 facilitates acetylation of proteins and RNA) our data indicated their mutual role of both proteins in DNA damage repair processes. Consistent with previous studies (Liu et al., 2016, Williams & Schumacher, 2016), we confirmed that NAT10 regulates the pool of DDR-related proteins, including p53 and DDB2, and likely participates in NER signaling. To this fact, original research (Liu et al., 2016) demonstrated that NAT10 acetylates p53 at lysine 120, stabilizing it via the MDM2 protein. This process activates p53-mediated cell cycle control following DNA damage. We tried to verify this data using AlphaFold 3 prediction tools and we observed a non-significant interaction between NAT10 and MDM2 proteins, while there is a pronounced interaction between ∼10-40 amino acids residues of p53 and ∼30-110 amino acids residues of the MDM2 protein (Supplementary Figure 7A and B). Performing AlphaFold 3 multimeric prediction, we observed non-significant changes for trimer NAT10-MDM2-p53 when compared with heterodimers NAT10-MDM2 or MDM2-p53 (Supplementary Figure 7A-C). Data only confirmed the interaction between p53 and MDM2 (Supplementary Figure 7B). Our experiments further showed a functional link between p53 and DDB2 (Figure 2E and F), consistent with earlier studies (Kannan, Amariglio et al., 2001, Wei, Wu et al., 2006). Importantly, in NAT10-deficient cells, the interaction between p53 and DDB2 was diminished (Figures 3D and E), suggesting that NAT10 indirectly influences DNA damage repair by modulating interactions of NER-related proteins, especially DDB2 and p53 (Figure 2I and Supplementary Figure 2A). To support this claim, did not find that the NAT10 acetyltransferase is directly recruited to UV-damaged chromatin (Svobodova Kovarikova et al., 2023).

Here, we also explore the impact of NAT10 depletion and UVC irradiation on genome-wide interactions and 3D-genome organization. The relationship between DNA damage and 3D-genome architecture remains an emerging area of study. Prior work (Sanders, Freeman et al., 2020) using Hi-C analysis revealed that radiation-induced changes in 3D-genome structure are cell-type specific, with stronger boundaries of TADs, observed in cells with functional ATM (the protein kinase Ataxia Telangiectasia Mutated). In our study, we observed significant differences in 3D-genome architecture when comparing NAT10 wild-type, NAT10-depleted, and UVC-irradiated cells, with pronounced changes in the organization of acrocentric chromosomes (Figure 5-7 and Supplementary Figures 3-5). For instance, in human acrocentric chromosomes, we observed different effects of UVC irradiation and NAT10 depletion. UVC irradiation uniformly changes the number of loop interactions in HSA13 (Figure 7A), while NAT10 depletion affects specific regions of HSA13 (Figure 7B). Moreover, UVC irradiation increases the number of chromatin loop interactions (Figure 7C), which might be connected with chromatin condensation. NAT10 depletion decreases the number of interactions (Figure 7C) supporting the role of NAT10 in the formation of nucleoli and chromatin decondensation. Using cLoops2 for the full-stack analysis of the 3D-chromatin interaction data, we identified differences in loop interactions across various conditions: NAT10 wt / NAT10 dn or NAT10 wt / NAT10 wt UVC or NAT10 dn / NAT10 dn UVC (Figure 5A-C, 6A-C). We have found that both NAT10 depletion, and to a lesser extent, UVC irradiation significantly impact chromatin loop formation in comparison with NAT10 wt cells. This claim is documented by differences in enrichment scores showing highly enriched chromatin loops and loops with reduced interaction properties (Figure 5A-C).

Overall, our findings highlight the complex interplay between NAT10, DDR-related proteins, and 3D-genome architecture in the context of DNA damage and repair. Thus, our study underscores the multifaceted role of NAT10 in coordinating chromatin organization, RNA acetylation, and DNA repair mechanisms, offering new perspectives on its function in genome stability. We showed that the 3D-chromatin organization of human acrocentric chromosomes undergoes significant alterations in cells depleted of the NAT10 acetyltransferase or exposed to UVC irradiation. This function is likely mediated through NAT10 interplay with key DNA damage response proteins, including p53 and DDB2, both of which are essential regulators of the Nucleotide Excision Repair (NER) pathway.

## Methods

### Cell cultivation and treatment

The human cervix adenocarcinoma (HeLa) NAT10 wild-type (wt) and NAT10 double null (dn) cells were a generous gift from Dr. Shalini Oberdoerffer, NCI NIH, USA (Arango et al., 2018, Arango, Sturgill et al., 2022). Both NAT10 wt and NAT10 dn cell lines were cultivated in DMEM (Dulbecco’s modified Eagle’s medium, Merck, Czech Republic) supplemented with 10% fetal calf serum (FCS, #FB-1003, BioTech, Czech Republic) and 2 mM L-glutamine (#25030081, Thermo Fisher Scientific, Czech Republic) without antibiotics.

The human non-small cell lung carcinoma cell line H1299 [p53(dn)] and H1299 [p53(wt), Tet-On system] were a generous gift from Dr. Marie Brázdová, Institute of Biophysics of the Czech Academy of Sciences, Czech Republic (Brazdova, Tichy et al., 2016, Rohaly, Chemnitz et al., 2005). The expression of p53 wt in the H1299 cell line was induced with 1 μg/ml tetracycline (#87128, Merck, Czech Republic) 24 hours before the harvesting. Cells were cultivated in DMEM medium supplemented with 10% FCS, 1 mM sodium pyruvate (#8636, Merck, Czech Republic), and the appropriate antibiotics (#XC-A4122, BioTech, Czech Republic). All cell lines were incubated in a humified atmosphere at 37 °C supplemented with 5% CO_2_.

Cells were grown to 75% confluence and then irradiated by the UVC lamp (Philips, Amsterdam, The Netherlands, model TUV 30 W T8, UVC 254 nm wavelength). Irradiation was performed for 10 min. In detail, 5 min after UVC irradiation, the cells were either fixed or harvested and proceeded with the Proximity Ligation Assay (PLA) or western blot as described below. The lamp distance from the sample was 60 cm (Svobodova Kovarikova et al., 2023, Svobodova Kovarikova, Stixova et al., 2020).

### Immunofluorescence staining and confocal microscopy

Immunofluorescence was modified following the approach by (Svobodova Kovarikova et al., 2020). The cells were fixed in 4% formaldehyde (FA; #AAJ19943K2, Fisher Scientific, USA) for 10 min at room temperature (RT), permeabilized with 0.2% Triton X-100 (#194854, MP Biomedicals, USA) for 8 min, and 0.1% saponin (#S7900, Merck, Germany) for 13 min. In the next step, the microscopic dishes were washed twice in phosphate buffer saline (PBS) for 15 min. We used 10% goat serum (#G9023, Merck, Germany) dissolved in 1x PBS-Triton X-100 (0.1%) as a blocking solution. The samples were incubated for one hour at room temperature and then washed in 1x PBS for 15 min. For immunofluorescence analysis, the following antibodies were used: anti-H3K9me3 (#ab8898, Abcam, UK), anti-H3K36me3 (#ab9050), anti-H3K79me1 (#ab2886), anti-H3K79me2 (#ab3596), anti-H3K79me3 (#ab2621), and anti-H3K9ac (#06-942, Merck, Czech Republic). The following secondary antibodies were used: Alexa Fluor 488-conjugated donkey anti-mouse (#A21202, Thermo Fisher Scientific, USA), Alexa Fluor 488-conjugated goat anti-rabbit (#ab150077, Abcam, UK), Alexa Fluor 594-conjugated goat anti-mouse (#A11032, ThermoFisher Scientific, USA), Alexa Fluor 594-conjugated goat anti-rabbit (#A11037), and goat anti-rabbit Cy5 (#ab6564, Abcam, UK). The secondary antibodies were diluted at 1:200, and the incubation time was 60 min at RT. A contour of cell nuclei (condensed chromatin) was visualized using 4′,6-diamidino-2-phenylindole (DAPI; #D9542, Merck, Germany), dissolved in Vectashield (#H-1000, Vector Laboratories, USA).

Fluorescence images were acquired using a Leica TCS SP8-X SMD confocal microscope (Leica Microsystem, Germany), equipped with oil CS2 objective (HC PL APO 63×, NA = 1.4). Image acquisition was performed using a white light laser (WLL; wavelengths of 470-670 nm in 1-nm increments) with the following parameters: 1024×1024 pixel resolution, 400 Hz, bidirectional mode, and zoom 2. As described above, we used Leica Application Suite (LAS X) software for immunofluorescence analysis.

### Proximity Ligation Assay (PLA)

The PLA technique was used to investigate protein-protein interactions in fixed cells. We studied the interactions between proteins p53-DDB2 and NAT10-DDB2 in NAT10 wt and NAT10 dn cell lines. Firstly, cells were grown to 75% confluence on coverslips (12 mm no. 1.5 round, #0112520, Paul Marienfeld GmbH., Germany) and 24 h after seeding, washed in PBS buffer, and fixed in 4% formaldehyde (FA; #AAJ19943K2, Fisher Scientific, USA) for 10 min at room temperature (RT). In the next step, cells were permeabilized with 0.2% Triton X-100 (#194854, MP Biomedicals, USA) for 8 min and 0.1% saponin (#S7900, Merck, Germany) for 13 min and then washed twice in phosphate buffer saline (PBS) for 15 min. After permeabilization, the non-specific binding sites were blocked by 1% Bovine Serum Albumin (BSA, #A2153, Meck, Czech Republic) dissolved in 1x PBS for 1 h at RT. As the final step, samples were washed for 15 min in PBS and incubated at 4 °C overnight with primary antibodies (dilution 1:100 in 1% BSA): anti-p53 (#sc-126, Santa Cruz Biotechnology Inc., USA), anti-NAT10 (#sc-271770) and anti-DDB2 (#sc-25368). PLA was performed according to the manufacturer’s protocol (Duolink® In Situ Orange Starter Kit, #DUO92102-1KT, Merck, Czech Republic). After overnight incubation with primary antibodies, samples were washed twice with 1x Wash Buffer A® for 5 minutes each at RT. In the next step, samples were incubated with the affinity-purified donkey anti-rabbit IgG Duolink in Situ PLA probe PLUS® (1:5, Sigma Aldrich) and affinity-purified donkey anti-mouse IgG Duolink in Situ PLA probe MINUS® (1:5, Sigma Aldrich), containing the secondary antibody conjugated with complementary oligonucleotides in a humid chamber at 37 °C for 1 hour. Incubation was followed by washing samples twice with 1x Wash Buffer A® for 5 minutes each at RT. After washing, samples were incubated in 1 unit/μl of T4 DNA ligase in diluted ligase buffer (1:5, Sigma Aldrich) for 30 min at 37 °C in the humid chamber. The ligase solution was washed twice with 1x Wash Buffer A® for 2 minutes at RT. The signals were then amplified using the amplification solution containing 5 units/μl of DNA polymerase in diluted polymerase buffer with orange fluorescent-labeled oligonucleotides (1:5, Sigma Aldrich) for 100 min at 37 °C in a humified chamber. A final round of washes was performed twice for 10 min in 1x Wash Buffer B®. The slides were then mounted using 5 μl of Duolink in situ mounting medium containing DAPI and visualized as described below. Appropriate controls were included, such as samples treated with only one primary antibody or no antibodies, to validate the specificity of the PLA signals. The amplified signals were visualized using a Leica TCS SP8-X SMD confocal microscope (Leica Microsystem, Germany) equipped with oil CS2 objective (HC PL APO 63×, NA = 1.4). Image acquisition was performed using a white light laser (WLL) with the following parameters: 1024×1024 pixel resolution, 400 Hz, bidirectional mode, and zoom 2. We used ImageJ software (NIH freeware, USA) to analyze the number of PLA signals (dots per nucleus). The statistical analysis was performed using GraphPad Prism 9 software (USA) and the nonparametric Mann–Whitney *U*-test. The asterisks in the figures represent statistical significance with a p-value ≤ 0.05 (*). At least 150 cells were compared within one experimental data set for each condition.

### Western blot

The western blot analysis was performed following (Legartova, Jugova et al., 2013). Protein levels were determined using a µQuant spectrophotometer (BioTek, USA) to ensure uniform concentrations of total proteins. Proteins were then separated by SDS polyacrylamide gel electrophoresis (SDS-PAGE) and transferred onto polyvinylidene difluoride (PVDF) membranes. Following this, the membranes were blocked with 5% non-fat dry milk for 2 hours and were incubated overnight at 4 °C with specific primary antibodies: anti-NAT10 (#sc-271770, Santa Cruz Biotechnology Inc., USA), anti-p53 (#sc-126), anti-XPC (#sc-30156); anti-XPA (#sc-28353), anti-DDB2 (#sc-25368), and anti-GAPDH (#60004-1, Proteintech, Germany). For the histone code studies we used the following antibodies: anti-H3K9me1 (#ab9045, Abcam, UK), anti-H3K9me2 (#ab2120), anti-H3K9me3 (#ab8898), anti-H3K36me3 (#ab9050), anti-H3K79me1 (#ab2886), anti-H3K79me2 (#ab3596), anti-H3K79me3 (#ab2621), anti-H3K9ac (#06-942, Merck, Czech Republic), anti-H4ac (#382160, Merck, Czech Republic), and anti-H3 (#ab1791). The next day, following several washes, the membranes were treated with secondary antibodies: goat anti-rabbit IgG (#AP307P, Merck, Czech Republic), rabbit anti-mouse IgG (#A9044), and goat anti-mouse IgG_1_ (#sc-2060, Santa Cruz Biotechnology Inc., USA), all of which were diluted 1:2000 in blocking solution. The density of the bands on the western blot was quantified using ImageJ software (NIH freeware, USA).

### Local laser microirradiation

For microirradiation experiments with UVA lasers (wavelengths 355 nm and 405 nm), cells were seeded on 35 mm gridded microscope dishes (#81,166, Ibidi, Fitchburg, WI, USA). At 50% confluence, they were sensitized with 10 μM 5-Bromo-2′-deoxyuridine (BrdU; #11296736001, Merck, Germany) for 16-18 hours. During microscopy, cells were maintained under optimal conditions in an incubation chamber (EMBL) at 37 °C with 5% CO₂. Irradiation was performed using a TCS SP5-X confocal microscope system (Leica, Germany) equipped with 355-nm and 405-nm lasers and a 63× oil objective (HCX PL APO, lambda blue) with a numerical aperture (NA) of 1.4. The microscope settings for local DNA damage induction were as follows: 355-nm laser power at 80 mW with 100% output, 16×16 pixel resolution, 10 Hz, bidirectional mode, line averaging of 32, and a zoom factor of 1.5×. Microirradiated cells were monitored at precise time points ranging from 2 minutes to 4 hours. Analysis was conducted on four biological replicates. Following the immunostaining procedure, locally microirradiated cells were found based on registered coordinates on gridded dishes. We examined the levels of DDB2 and γH2AX [phospho-histone H2A.X (Ser139)]. LEICA LAS X software was used for image acquisition and fluorescence intensity (FI) analysis.

### Hi-C methodology and statistical analysis

Hi-C (High-throughput Chromosome Conformation Capture) analysis was performed in the core facility of the Active Motif company focusing on epigenetic research. Using this technique, we studied the three-dimensional architecture of the human genome in HeLa tumor cells. By capturing and sequencing DNA fragments, that are close to the cell nucleus, Hi-C analysis of non-irradiated and UVC-irradiated NAT10 wt and NAT10 dn cells generated a comprehensive map of chromosomal interactions (3D-genome interactions). For such analysis, HeLa cells were cultivated till the confluence of 70-80 % and after that, cells were irradiated by a UVC lamp, as described above. The pellet of 10^6^ non-irradiated and UVC-irradiated NAT10 wt and NAT10 dn cells were harvested and frozen at −80 °C. These samples were processed, via Active Motif service, in the NIH core facility interested in Hi-C analysis. The samples were prepared for library generation using the Arima-HiC Kit sample preparation protocol (A510008, Arima Genomics, USA). Following the manufacturer’s instructions, the samples were crosslinked and lysed, followed by restriction enzyme digestion, biotinylation, and ligation. The ligated DNA was then purified and sheared using the PIXUL™ Multi-Sample Sonicator (PIXUL, #53130, Active Motif, USA). The sheared DNA was subsequently used for library generation with the 2S™ Plus DNA Library Kit (#10009877, Integrated DNA Technologies, IDT, USA). The samples were indexed using the 2S™ MID Adapter Set A and/or B (#10009900/#10009901, IDT, USA). Library amplification was performed using the KAPA HiFi HotStart Library Amplification Kit with Primer Mix (#07958978001, Roche Sequencing Solutions, RSS, USA). Libraries were quantified using a Qubit Fluorometer (ThermoFisher Scientific, USA) and a TapeStation (Agilent, USA), and sequenced with paired-end 150 base pair reads (PE150) on the Illumina NovaSeq platform (Novogene Corporation, USA). The processing and statistical analysis were performed according to (Servant, Varoquaux et al., 2015). The Hi-C data analysis begins with the alignment of raw Hi-C reads to a reference human genome (hg38) to identify interacting DNA fragments. Following alignment, the data undergoes a series of quality control (QC) checks to remove artifacts and ensure the reliability of detected interactions. Next, interaction contact matrices are constructed, representing the frequency of contacts between different genomic regions. These matrices are then normalized to correct for biases inherent in the experimental procedure. In detail, raw genome maps are shown in resolution 1Mb/bin (Supplementary Figures 3-5, panels A and C), and contact maps at resolution 360 Kb/bin are shown (Supplementary Figure 3, 4 in panels B, D, and 5B). Difference maps, representing changes between two datasets, show an increased number of interactions (red) and a reduced number of interactions (blue) (Supplementary Figure 3, 4 in panels B, D, and 5B, red-blue panels). Differential maps, representing aggregation features of chromatin loops by mathematic approaches, are shown in Figure 5A-C. Advanced statistical and computational methods were applied to identify significant chromatin interactions and structural features, such as differential loops, and A/B compartments. The A/B compartments were estimated using eigenvector analysis of the genome contact maps. Cooltools package (https://github.com/open2c/cooltools) was used to determine compartments from Hi-C data. Visualization tools and further analyses were used to interpret the results, linking chromatin architecture to gene regulation, epigenetic modifications, and cellular functions. The nf-core/hic pipeline was applied to process the Hi-C data. Generally, differential loop analysis describes the aggregated features of common (unspecific) chromatin loops and unique (conditional specific) chromatin loops between two conditions using cLoops2 tool (Cao, Liu et al., 2022). Here, we especially focused on the 3D-nuclear architecture of acrocentric chromosomes, because NAT10, a main protein of our interest, mainly occupies the compartment of the nucleolus (Figure 1A). It is also known that p-arms of acrocentric chromosomes (HSA13-15, 21, and 22) mainly form nucleoli, but sequences elsewhere on these chromosomes can associate with nucleolar regions (van Sluis, van Vuuren et al., 2020). Therefore, regions of Hi-C matrices representing individual acrocentric chromosomes were visualized in HiGlass software (https://higlass.io/app) at the resolution of 360 kilobases per bin. The visualizations (Figure 7A and B and Supplementary Figures 3-5) show the difference between experimental datasets. We compared differences between non-irradiated NAT10 wt HeLa cells and their UVC-irradiated counterpart, or non-irradiated NAT10 wt HeLa cells and non-irradiated NAT10 dn HeLa cells; or non-irradiated NAT10 dn HeLa cells and UVC-irradiated NAT10 dn HeLa cells. Furthermore, we compared the decay of interaction counts with genomic distance across all experimental datasets using HiCExplorer’s hicPlotDistVsCounts tool https://hicexplorer.readthedocs.io/en/latest/content/tools/hicPlotDi <u>stVsCounts.html</u>).

### AlphaFold 3 dimer and multimer protein predictions

AlphaFold 3 predictions were performed for proteins whose sequences are available at https://www.uniprot.org. Graphs show the pLDDT score and pTM. pLDDT is a per-atom confidence estimate on a 0-100 scale where a higher value indicates higher confidence. pLDDT aims to predict a modified LDDT score that only considers distances to polymers. The predicted template modeling (pTM) score is derived from a measure called the template modeling (TM) score. This measures the accuracy of the entire structure (Xu & Zhang, 2010, Zhang & Skolnick, 2004) pTM score above 0.5 means the overall predicted fold for the complex might be similar to the true structure. We also show panels in Supplementary Figure 1 that were generated using ChimeraX 1.9. Regions of low prediction confidence were not hidden. We predicted a degree of interactions between the NAT10 acetyltransferase and 18S rRNA (NR_003286.4, Homo sapiens RNA, 18S ribosomal N5 (RNA18SN5), ribosomal RNA, 1869 bp, gene) or NAT10 with the following proteins:

RNA18SN5: https://www.ncbi.nlm.nih.gov/nuccore/NR_003286

NAT10: https://www.uniprot.org/uniprotkb/Q9H0A0/entry,

p53: https://www.uniprot.org/uniprotkb/P04637/entry,

DDB1: https://www.uniprot.org/uniprotkb/Q16531/entry

DDB2: https://www.uniprot.org/uniprotkb/Q92466/entry

MDM2: https://www.uniprot.org/uniprotkb/Q00987/entry

## Data availability

The source data from Hi-C and western blots or AlphaFold 3 prediction will be shared or seen at https://www.ibp.cz/en/research/departments/cellular-biology-and-epigenetics/open-data. Alternatively, data are on request to the corresponding author.

## Author contributions

Eva Bártová: Conceptualization, writing original draft, writing review & editing, microscopy of Figures 1A, 4C, D, and 9G-L. Magdalena Skalníková: Formal analysis, methodology, writing review & editing, sample preparation for Hi-C, and Hi-C analysis. Lenka Stixová: Formal analysis, methodology, validation, editing, data on XPC in Figure 2A and microirradiation shown in Figure 4E, sample preparation for Hi-C. Vlastimil Tichý: Formal analysis, methodology, editing, Hi-C technical support. Filip Opálený and Jan Byška: analysis of the Hi-C data in Figure 7 and Supplementary Figures 3-5. Soňa Legartová: Formal analysis, methodology, and validation, SL is responsible for data in Figures 2A-F, 3B-F, and 8C-F. Tomáš Brom: AlphaFold 3 predictions.

## Acknowledgments

Many thanks to Dr. Petr Fajkus for his help with the illustration created using the BioRender software. Also, many thanks to Dr. Shuxiong Wang, from the Active Motif company, for helping with Hi-C data interpretation. Funded by the Institute of Biophysics of the Czech Academy of Sciences (IC: 68081707). The paper was supported by the Czech Science Foundation, grant no. GA23-05651S, and the Ministry of Education Youth and Sports of the Czech Republic, LUAUS25085.

## Disclosure and competing interest statement

The authors declare no competing interests.

## Supplementary Material

**Supplementary Figure 1.**
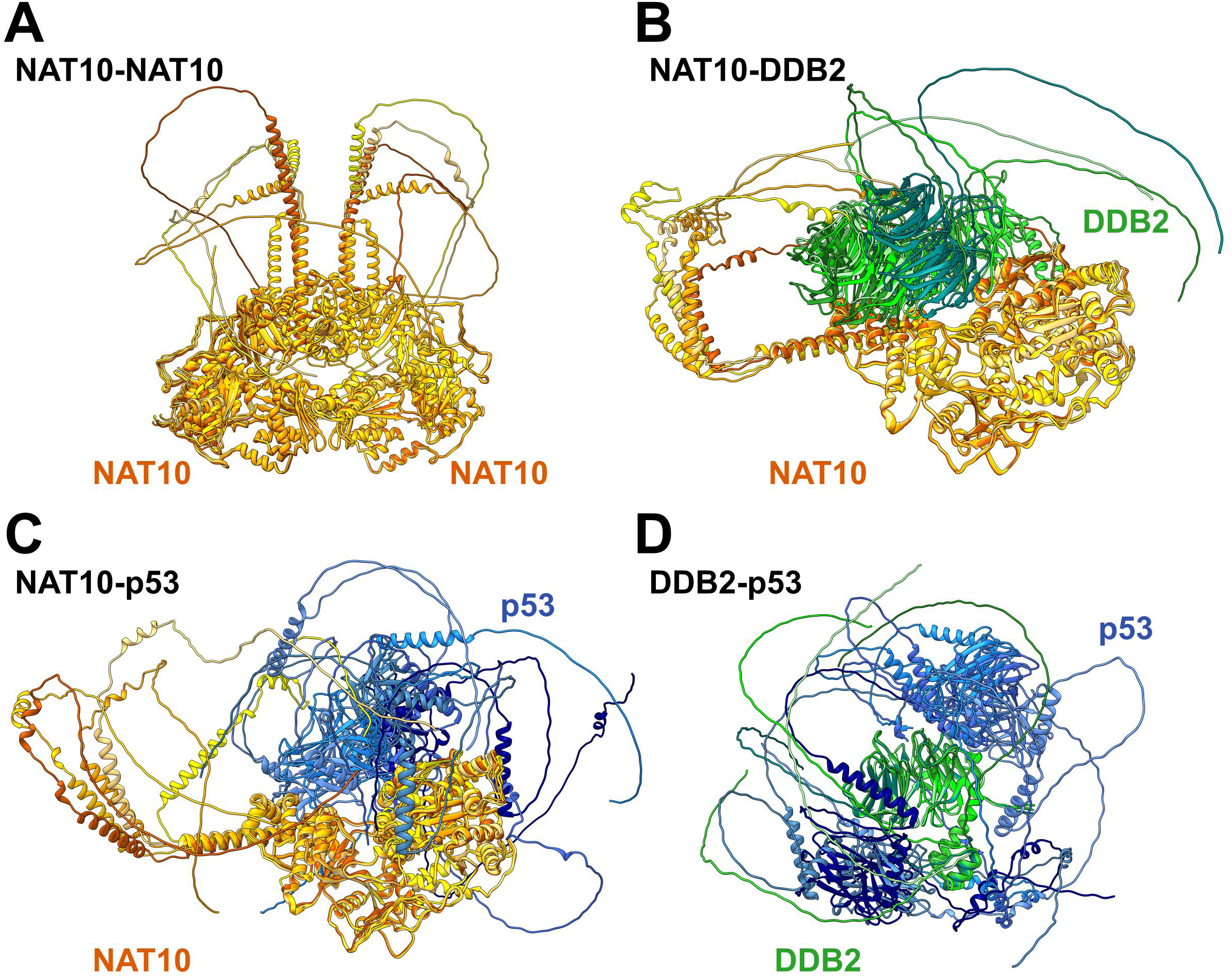
Data from AlphaFold 3, predicting **(A)** NAT10 dimerization and the degree of interaction between **(B)** NAT10-DDB2, **(C)** NAT10-p53, and **(D)** DDB2-p53, were imported to ChimeraX 1.9 software showing individual proteins. NAT10 is highlighted by orange, DDB2 is green, p53 is blue colour. The source data are shown at https://www.ibp.cz/en/research/departments/cellular-biology-and-epigenetics/open-data. This raw data from AlphaFold 3 can be uploaded to ChimeraX 1.9 for more detailed analysis.

**Supplementary Figure 2.**
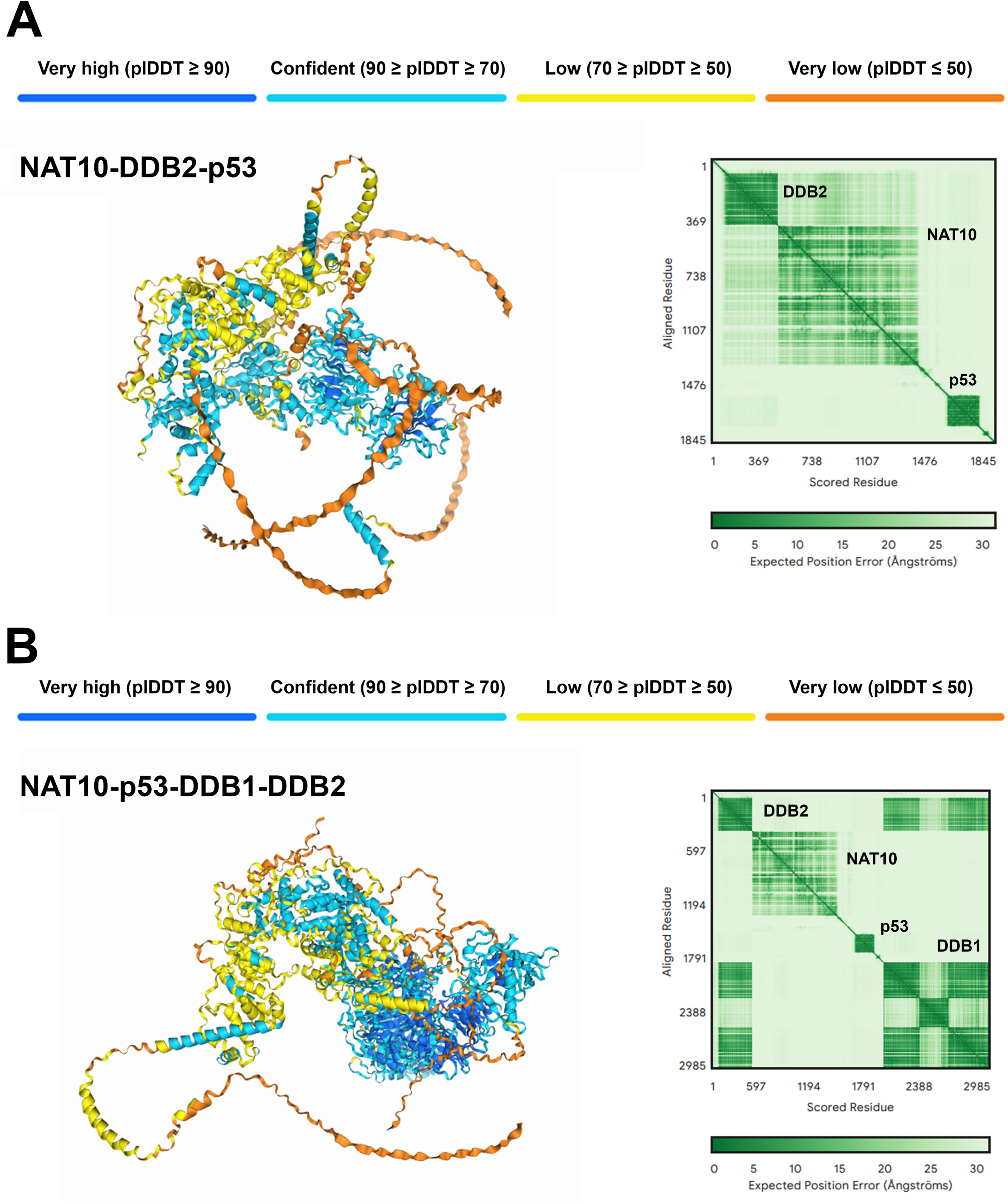
Panels **(A, B)** show AlphaFold 3 multimer prediction of human NAT10 interaction with **(A)** NAT10 and p53 and DDB2 proteins (pTM = 0.48) or **(B)** NAT10 and p53 and DDB1-DDB2 protein complex complex (pTM = 0.53).

**Supplementary Figure 3.**
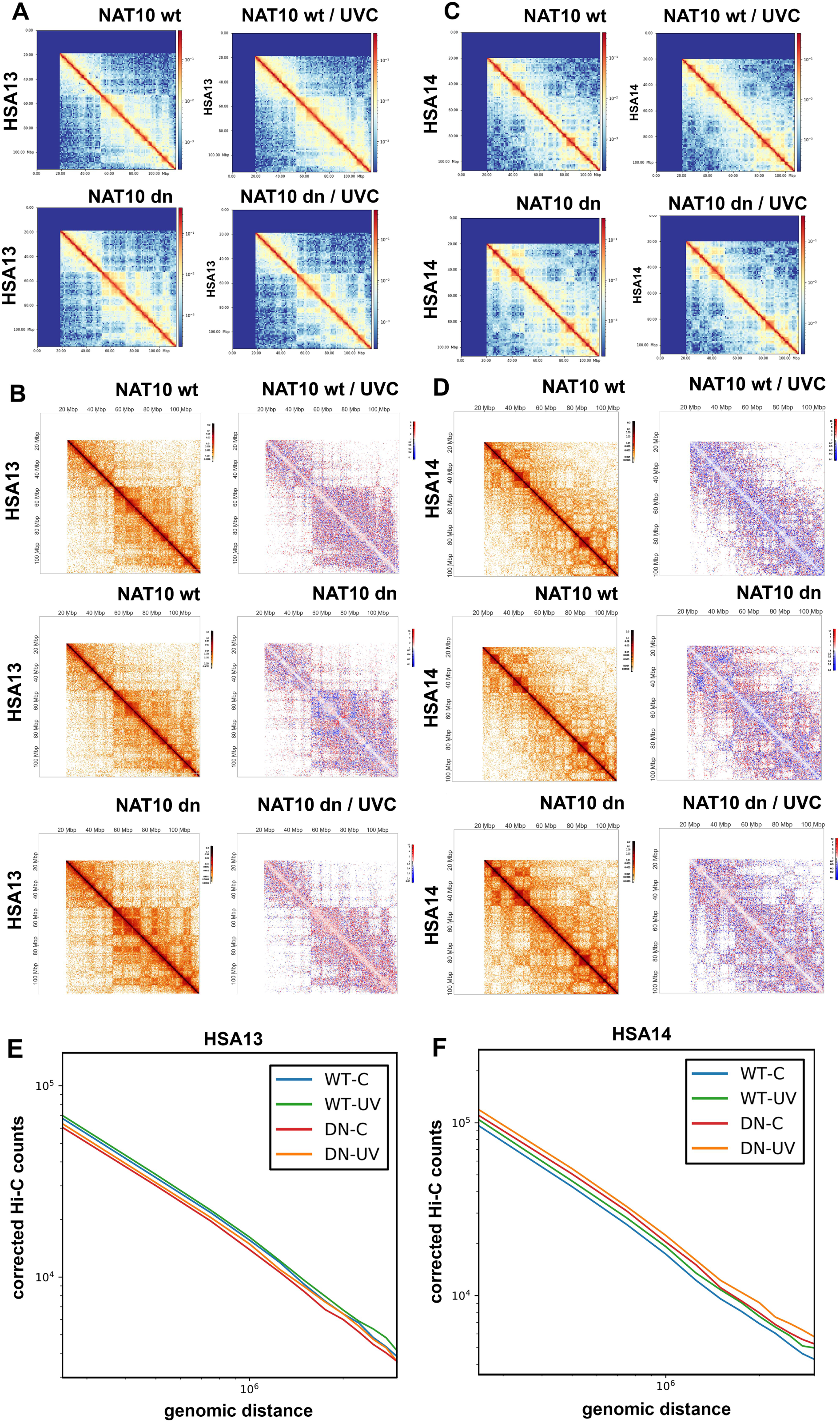
Changes in 3D-genome nuclear architecture of human acrocentric chromosomes 13 and 14 caused by NAT10 deficiency and UVC irradiation in HeLa cells. Genome-wide contact maps are shown for acrocentric chromosomes **(A)** HSA13 and **(C)** HSA14 in 1 Mb per bin. Panels **(B)** and **(D)** show regions of Hi-C matrices corresponding to individual acrocentric chromosomes that were visualized using the HiGlass software at a resolution of 360 Kb per bin (left panel in B and D). The visualization shows difference maps between the control (NAT10 wt or NAT10 dn) and comparative datasets (NAT10 wt / UVC; NAT10 dn or NAT10 dn / UVC) (red-blue right panels in B and D). The missing data in panels B and D are shown in white color. The following panels show global changes in average genomic interactions over all genomic distances of HSA13 **(E)** and HSA14 **(F)** studied in non-irradiated and UVC-irradiated NAT10 dn and NAT10 wt cells.

**Supplementary Figure 4.**
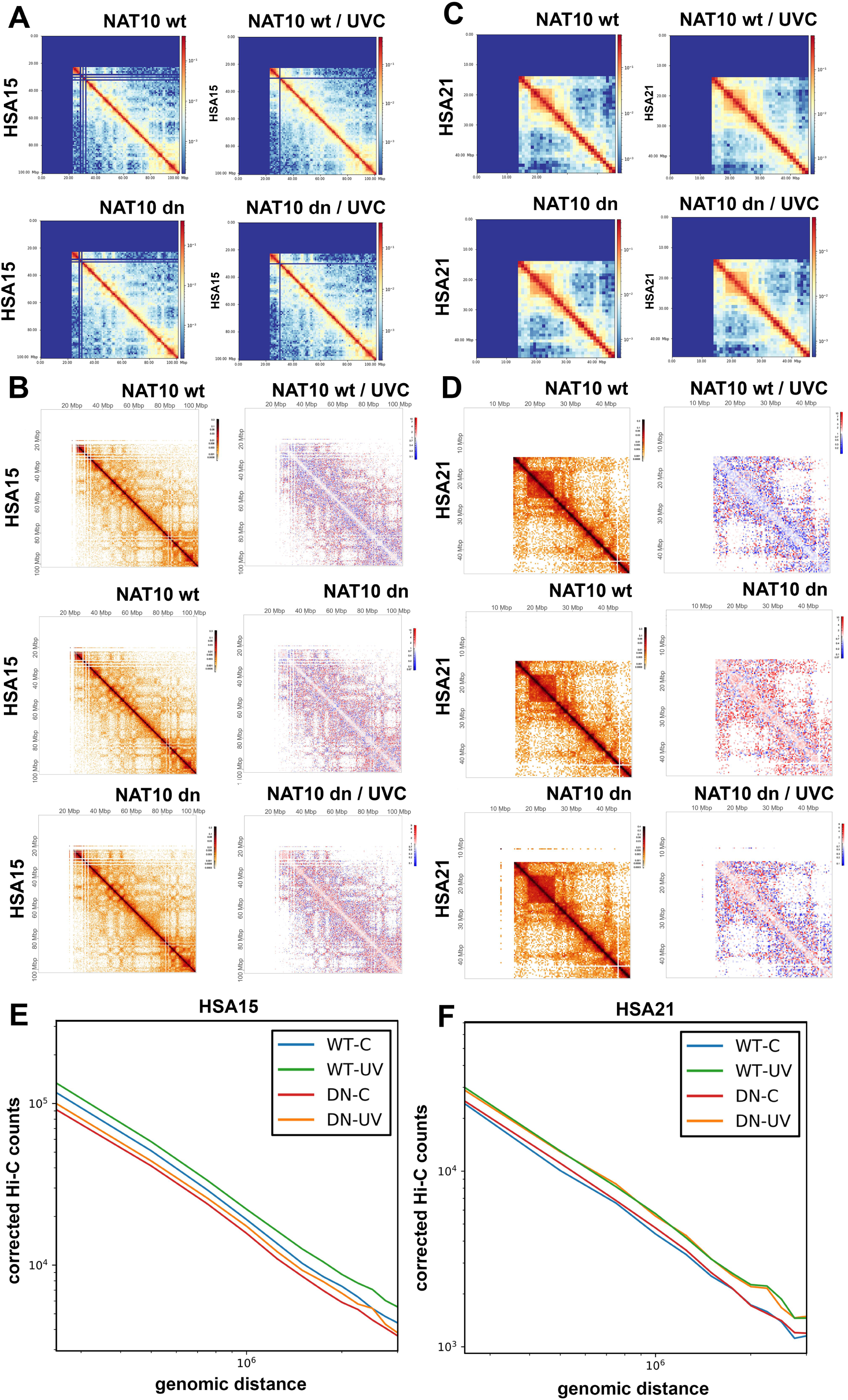
Changes in 3D-genome nuclear architecture of human acrocentric chromosomes 15 and 21 caused by NAT10 deficiency and UVC irradiation in HeLa cells. Genome-wide contact maps are shown for acrocentric chromosomes **(A)** HSA15 and **(C)** HSA21 in 1 Mb per bin. Panels **(B)** and **(D)** show regions of Hi-C matrices corresponding to HSA15 and HSA21 that were visualized using the HiGlass software at a resolution of 360 Kb per bin (left panels in B and D). The visualization shows difference maps between the control (NAT10 wt or NAT10 dn) and comparative datasets (NAT10 wt / UVC; NAT10 dn or NAT10 dn / UVC) (red-blue right panels in B and D). The missing data in panels B and D are shown in white color. The following panels show global changes in average genomic interactions over all genomic distances of HSA15 **(E)** and HSA21 **(F)** studied in non-irradiated and UVC-irradiated NAT10 dn and NAT10 wt cells.

**Supplementary Figure 5.**
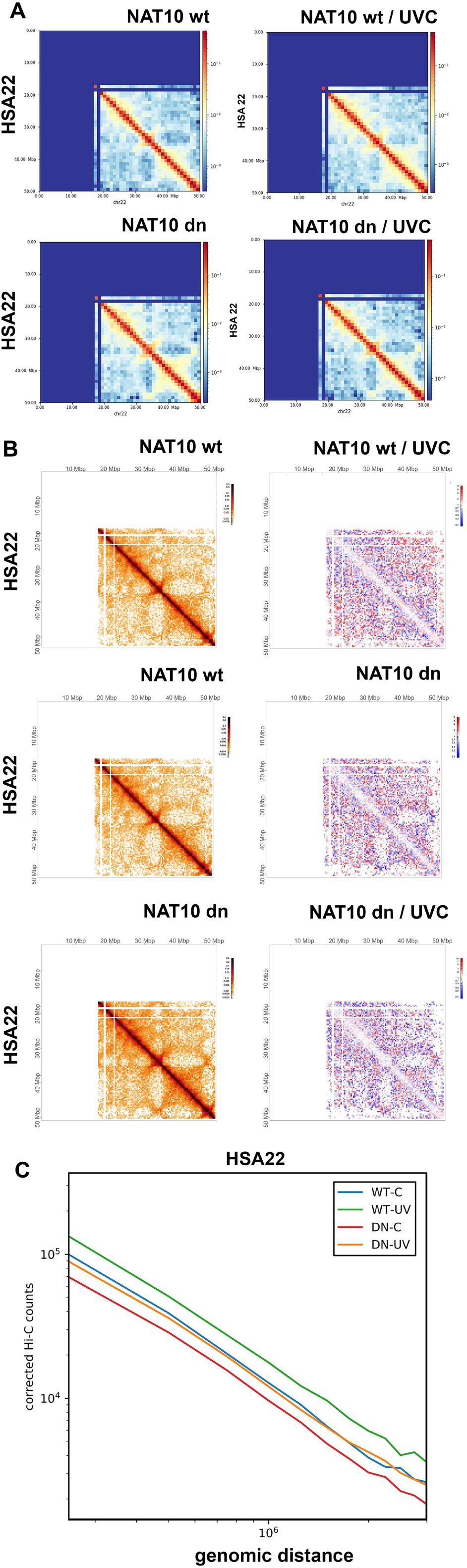
Changes in 3D-genome nuclear architecture of human acrocentric chromosome 22 caused by NAT10 deficiency and UVC irradiation in HeLa cells. Panel **(A)** shows contact maps of HSA22 analyzed in resolution 1 Mb per bin, studied in non-irradiated and UVC-irradiated NAT10 wt and NAT10 dn HeLa cells. Panel **(B)** documents regions of Hi-C matrices corresponding to individual acrocentric chromosome 22 visualized using the HiGlass software at a resolution of 360 Kb per bin (left panels in B and D). The visualization shows differences between the control (NAT10 wt or NAT10 dn) and comparative datasets (NAT10 wt / UVC; NAT10 dn or NAT10 dn / UVC) (red-blue right panels in B). The missing data in panel B are shown in white color. Panels **(C)** show global changes in average genomic interactions and overall genomic distances of HSA22 studied in non-irradiated and UVC-irradiated NAT10 dn and NAT10 wt cells. Hi-C data were also analyzed as an average over **(D)** all acrocentric chromosomes and compared with the average of **(E)** all human chromosomes and **(F)** all chromosomes without acrocentric ones.

**Supplementary Figure 6.**
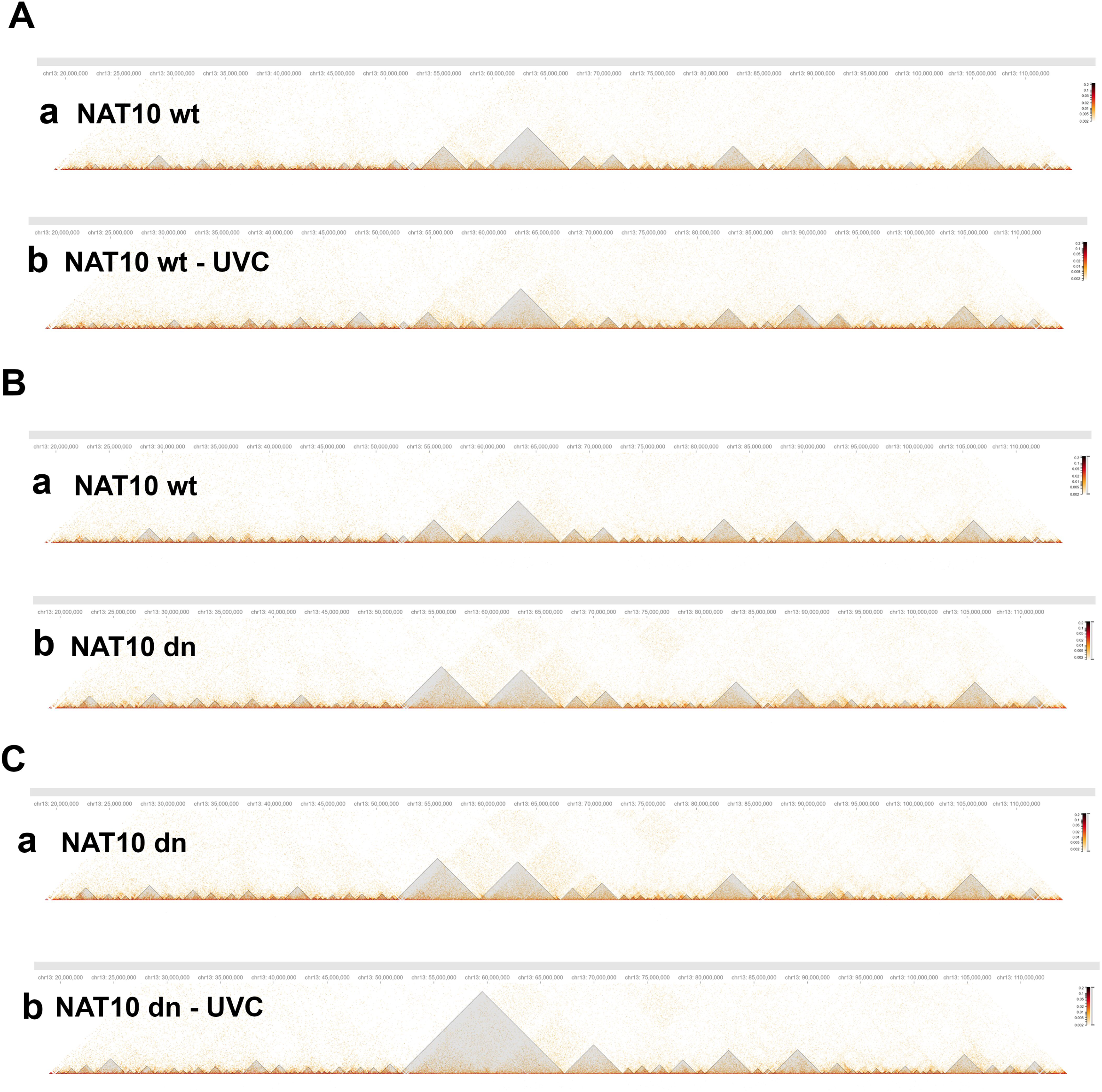
TADs are shown in resolution 40 kb per bin for human acrocentric chromosome 13. TADs were studied in non-irradiated and UVC-irradiated NAT10 wt and dn HeLa cells.

**Supplementary Figure 7.**
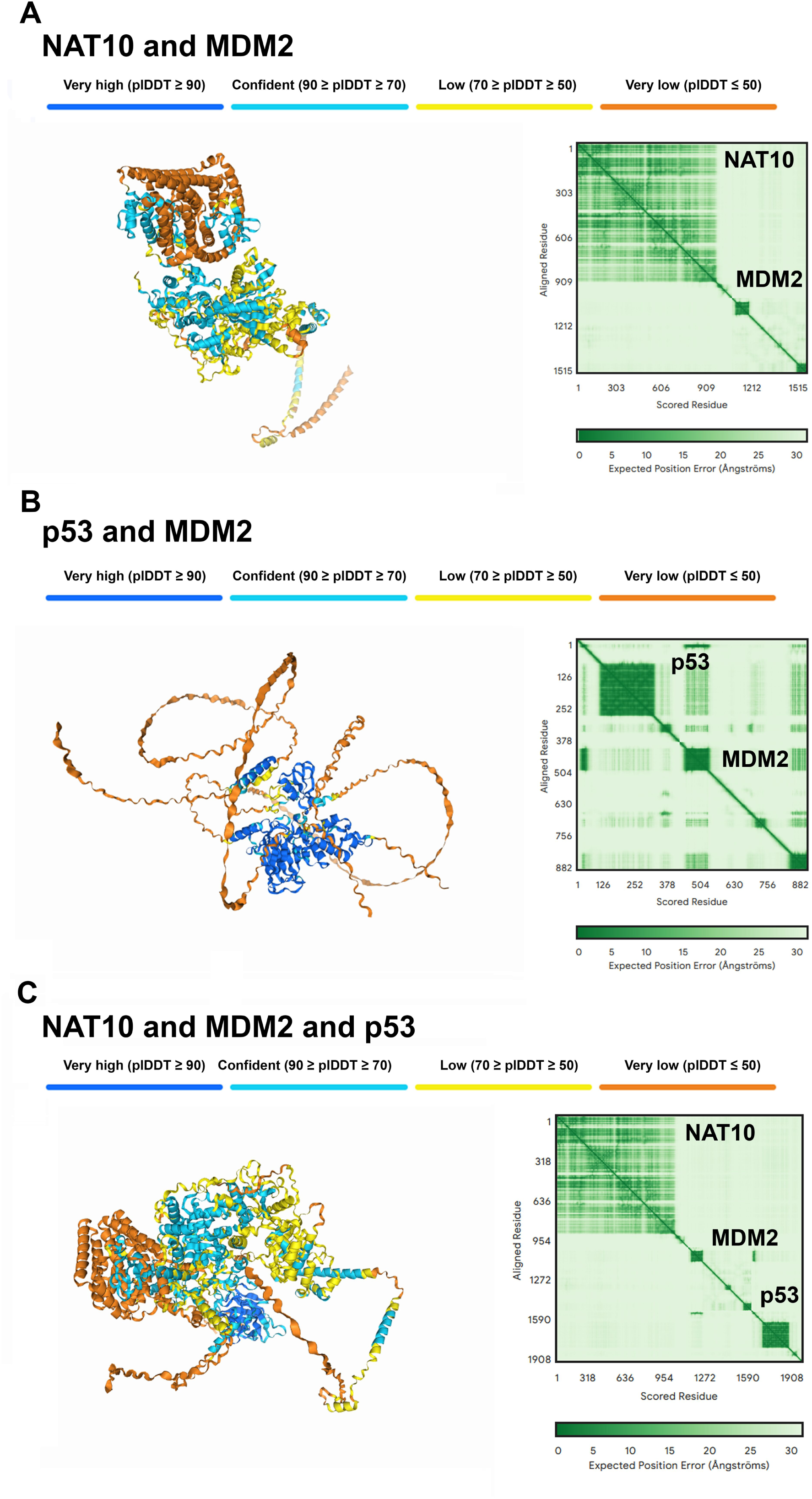
Results from AlphaFold 3 predictions of **(A)** NAT10 interaction with MDM2 (pTM = 0.49) and **(B)** MDM2 DNA repair-related protein interaction with p53 (pTM = 0.35). **(C)** Interaction properties of NAT10-MDM2-p53 trimer (pTM = 0.45) For source data, see https://www.ibp.cz/en/research/departments/cellular-biology-and-epigenetics/open-data.

**Supplementary Table 1.** Statistics of TADs on individual chromosomes: number of TADs (#), average size (Avg), and size of the largest TAD (kb).

## References

Arango D, Sturgill D, Alhusaini N, Dillman AA, Sweet TJ, Hanson G, Hosogane M, Sinclair WR, Nanan KK, Mandler MD, Fox SD, Zengeya TT, Andresson T, Meier JL, Coller J, Oberdoerffer S (2018) Acetylation of Cytidine in mRNA Promotes Translation Efficiency. Cell 175: 1872–1886 e24

Arango D, Sturgill D, Yang R, Kanai T, Bauer P, Roy J, Wang Z, Hosogane M, Schiffers S, Oberdoerffer S (2022) Direct epitranscriptomic regulation of mammalian translation initiation through N4-acetylcytidine. Mol Cell 82: 2797–2814 e11

Brazdova M, Tichy V, Helma R, Bazantova P, Polaskova A, Krejci A, Petr M, Navratilova L, Ticha O, Nejedly K, Bennink ML, Subramaniam V, Babkova Z, Martinek T, Lexa M, Adamik M (2016) p53 Specifically Binds Triplex DNA In Vitro and in Cells. PLoS One 11: e0167439

Cai S, Liu X, Zhang C, Xing B, Du X (2017) Autoacetylation of NAT10 is critical for its function in rRNA transcription activation. Biochem Biophys Res Commun 483: 624–629

Cao Y, Liu S, Ren G, Tang Q, Zhao K (2022) cLoops2: a full-stack comprehensive analytical tool for chromatin interactions. Nucleic Acids Res 50: 57–71

Capuozzo M, Santorsola M, Bocchetti M, Perri F, Cascella M, Granata V, Celotto V, Gualillo O, Cossu AM, Nasti G, Caraglia M, Ottaiano A (2022) p53: From Fundamental Biology to Clinical Applications in Cancer. Biology (Basel) 11

Chi YH, Haller K, Peloponese JM, Jr., Jeang KT (2007) Histone acetyltransferase hALP and nuclear membrane protein hsSUN1 function in de-condensation of mitotic chromosomes. J Biol Chem 282: 27447–27458

Ikeuchi Y, Kitahara K, Suzuki T (2008) The RNA acetyltransferase driven by ATP hydrolysis synthesizes N4-acetylcytidine of tRNA anticodon. EMBO J 27: 2194–203

Ito S, Horikawa S, Suzuki T, Kawauchi H, Tanaka Y, Suzuki T, Suzuki T (2014) Human NAT10 is an ATP-dependent RNA acetyltransferase responsible for N4-acetylcytidine formation in 18 S ribosomal RNA (rRNA). J Biol Chem 289: 35724–30

Kannan K, Amariglio N, Rechavi G, Jakob-Hirsch J, Kela I, Kaminski N, Getz G, Domany E, Givol D (2001) DNA microarrays identification of primary and secondary target genes regulated by p53. Oncogene 20: 2225–34

Lee JM, Hammaren HM, Savitski MM, Baek SH (2023) Control of protein stability by post-translational modifications. Nat Commun 14: 201

Legartova S, Jugova A, Stixova L, Kozubek S, Fojtova M, Zdrahal Z, Lochmanova G, Bartova E (2013) Epigenetic aspects of HP1 exchange kinetics in apoptotic chromatin. Biochimie 95: 167–79

Liu X CS, Zhang Ch, Liu Z, Luo J, Xing B, Du X. (2018) Deacetylation of NAT10 by Sirt1 promotes the transition from rRNA biogenesis to autophagy upon energy stress. Nucleic Acids Research, 2018, Vol 46, No 18 9601–96 46

Liu X, Tan Y, Zhang C, Zhang Y, Zhang L, Ren P, Deng H, Luo J, Ke Y, Du X (2016) NAT10 regulates p53 activation through acetylating p53 at K120 and ubiquitinating Mdm2. EMBO Rep 17: 349–66

Lv J, Liu H, Wang Q, Tang Z, Hou L, Zhang B (2003) Molecular cloning of a novel human gene encoding histone acetyltransferase-like protein involved in transcriptional activation of hTERT. Biochem Biophys Res Commun 311: 506–13

Rohaly G, Chemnitz J, Dehde S, Nunez AM, Heukeshoven J, Deppert W, Dornreiter I (2005) A novel human p53 isoform is an essential element of the ATR-intra-S phase checkpoint. Cell 122: 21–32

Sanders JT, Freeman TF, Xu Y, Golloshi R, Stallard MA, Hill AM, San Martin R, Balajee AS, McCord RP (2020) Radiation-induced DNA damage and repair effects on 3D genome organization. Nat Commun 11: 6178

Scrima A, Konickova R, Czyzewski BK, Kawasaki Y, Jeffrey PD, Groisman R, Nakatani Y, Iwai S, Pavletich NP, Thoma NH (2008) Structural basis of UV DNA-damage recognition by the DDB1-DDB2 complex. Cell 135: 1213–23

Servant N, Varoquaux N, Lajoie BR, Viara E, Chen CJ, Vert JP, Heard E, Dekker J, Barillot E (2015) HiC-Pro: an optimized and flexible pipeline for Hi-C data processing. Genome Biol 16: 259

Sharma S, Langhendries JL, Watzinger P, Kotter P, Entian KD, Lafontaine DL (2015) Yeast Kre33 and human NAT10 are conserved 18S rRNA cytosine acetyltransferases that modify tRNAs assisted by the adaptor Tan1/THUMPD1. Nucleic Acids Res 43: 2242–58

Shen Q, Zheng X, McNutt MA, Guang L, Sun Y, Wang J, Gong Y, Hou L, Zhang B (2009) NAT10, a nucleolar protein, localizes to the midbody and regulates cytokinesis and acetylation of microtubules. Exp Cell Res 315: 1653–67

Stoyanova T, Roy N, Kopanja D, Raychaudhuri P, Bagchi S (2009) DDB2 (damaged-DNA binding protein 2) in nucleotide excision repair and DNA damage response. Cell Cycle 8: 4067–71

Svobodova Kovarikova A, Stixova L, Kovarik A, Bartova E (2023) PARP-dependent and NAT10-independent acetylation of N4-cytidine in RNA appears in UV-damaged chromatin. Epigenetics Chromatin 16: 26

Svobodova Kovarikova A, Stixova L, Kovarik A, Komurkova D, Legartova S, Fagherazzi P, Bartova E (2020) N(6)-Adenosine Methylation in RNA and a Reduced m(3)G/TMG Level in Non-Coding RNAs Appear at Microirradiation-Induced DNA Lesions. Cells 9

Tan Y, Zheng J, Liu X, Lu M, Zhang C, Xing B, Du X (2018) Loss of nucleolar localization of NAT10 promotes cell migration and invasion in hepatocellular carcinoma. Biochem Biophys Res Commun 499: 1032–1038

Tang J, Chu G (2002) Xeroderma pigmentosum complementation group E and UV-damaged DNA-binding protein. DNA Repair (Amst) 1: 601–16

van Sluis M, van Vuuren C, Mangan H, McStay B (2020) NORs on human acrocentric chromosome p-arms are active by default and can associate with nucleoli independently of rDNA. Proc Natl Acad Sci U S A 117: 10368–10377

von Eyss B, Maaskola J, Memczak S, Mollmann K, Schuetz A, Loddenkemper C, Tanh MD, Otto A, Muegge K, Heinemann U, Rajewsky N, Ziebold U (2012) The SNF2-like helicase HELLS mediates E2F3-dependent transcription and cellular transformation. EMBO J 31: 972–85

Wei CL, Wu Q, Vega VB, Chiu KP, Ng P, Zhang T, Shahab A, Yong HC, Fu Y, Weng Z, Liu J, Zhao XD, Chew JL, Lee YL, Kuznetsov VA, Sung WK, Miller LD, Lim B, Liu ET, Yu Q et al. (2006) A global map of p53 transcription-factor binding sites in the human genome. Cell 124: 207–19

Williams AB, Schumacher B (2016) p53 in the DNA-Damage-Repair Process. Cold Spring Harb Perspect Med 6

Wittschieben BO, Wood RD (2003) DDB complexities. DNA Repair (Amst) 2: 1065–9

Xu J, Zhang Y (2010) How significant is a protein structure similarity with TM-score = 0.5? Bioinformatics 26: 889–95

Yang Z. WE, Cui Y.H., Li H., He Y-Y. (2023) NAT10 regulates the repair of UVB-induced DNA damage and tumorigenicity. Toxicology and Applied Pharmacology 477

Zhang XP, Liu F, Wang W (2011) Two-phase dynamics of p53 in the DNA damage response. Proc Natl Acad Sci U S A 108: 8990–5

Zhang Y, Skolnick J (2004) Scoring function for automated assessment of protein structure template quality. Proteins 57: 702–10

Zhou M, Gamage ST, Tran KA, Bartee D, Wei X, Yin B, Berger S, Meier JL, Marmorstein R (2024) Molecular Basis for RNA Cytidine Acetylation by NAT10. bioRxiv doi: 10.1101/2024.03.27.587050

